# Prefrontal cortex exhibits multi-dimensional dynamic encoding during decision-making

**DOI:** 10.1101/808584

**Authors:** Mikio C. Aoi, Valerio Mante, Jonathan W. Pillow

## Abstract

Recent work has suggested that prefrontal cortex (PFC) plays a key role in context-dependent perceptual decision-making. Here we investigate population-level coding of decision variables in monkey PFC using a new method for identifying task-relevant dimensions of neural activity. Our analyses reveal that, in contrast to one-dimensional attractor models, PFC has a multi-dimensional code for decisions, context, and relevant as well as irrelevant sensory information. Moreover, these representations evolve in time, with an early linear accumulation phase followed by a phase with rotational dynamics. We identify the dimensions of neural activity associated with these phases, and show that they are not the product of distinct populations, but of a single population with broad tuning characteristics. Finally, we use model-based decoding to show that the transition from linear to rotational dynamics coincides with a sustained plateau in decoding accuracy, revealing that rotational dynamics in PFC preserve sensory as well as choice information for the duration of the stimulus integration period.

## Introduction

A large body of work has aimed to identify the precise computational roles of various brain regions during perceptual decision-making ^1–8^. Recent interest has centered on prefrontal cortex (PFC), which has been shown to carry a wide range of sensory, cognitive, and motor signals relevant for integrating sensory information and making decisions ^1;4;6;7;9–12^. A major barrier to understanding PFC’s functional role, however, is that PFC neurons exhibit mixed selectivity, characterized by heterogeneous tuning to multiple task variables ^13^. The idiosyncratic single-neuron responses observed in PFC make it difficult to gain insight into the population-level representation of different sensory and cognitive variables. ^14–16^.

Here we analyze the population-level representation of information in PFC using *model-based targeted dimensionality reduction (mTDR)*, a general method for identifying the dimensions of population activity that encode information about different task variables over time. We applied this method to data from a context-dependent perceptual decision-making task ^1^, in which a context cue determined what kind of sensory information (color or motion) should be used for making a binary decision (Fig. 2a,b). In contrast to previous findings, our analysis revealed that the encoding of decisions, context, and relevant as well as irrelevant stimulus variables exhibited rotational dynamics in a multi-dimensional subspace, involving modulation of two or more orthogonal neural activity patterns over time.

**Figure 1:**
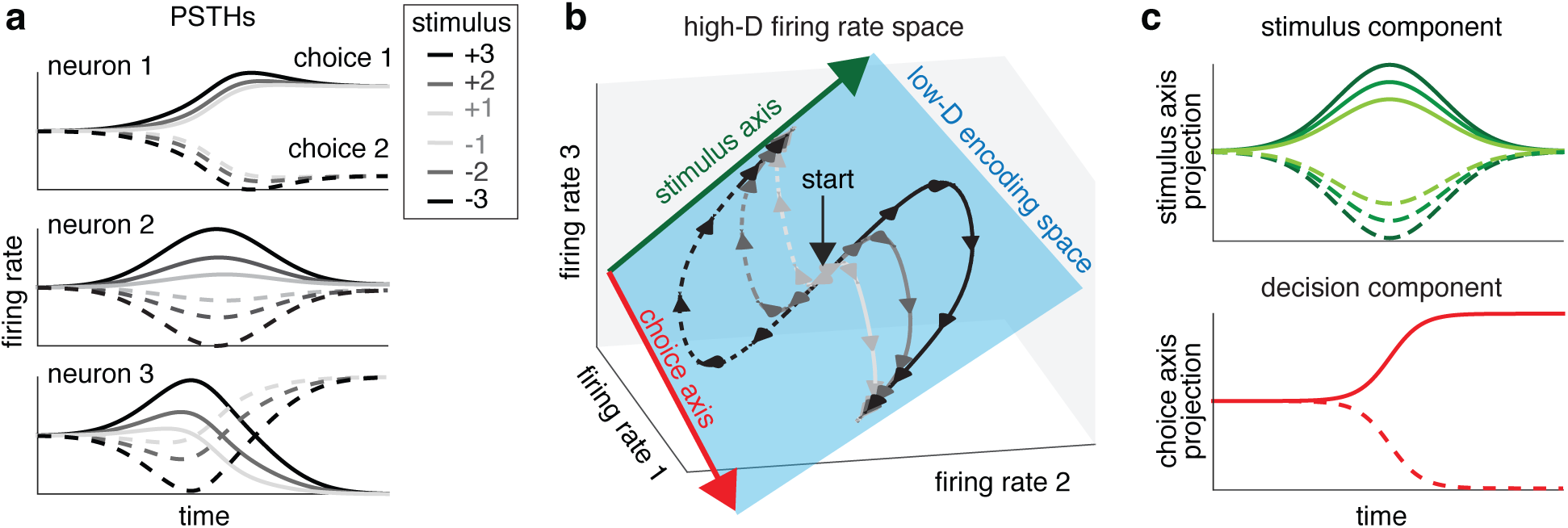
Schematic illustrating low-dimensional population-level encoding in a binary sensory decision-making task. **(a)**. Conditional PSTHs for three neurons that exhibit mixed selectivity to a stimulus variable (taking on six different values) and a choice variable (taking on two values). **(b)** Modulations of the PSTHs by the task variables span a 2-dimensional “encoding subspace”, which is low-dimensional relative to the 3-dimensional space of firing rates. In this case, a 1D stimulus-encoding subspace (green arrow) captures all information about the stimulus value, while a 1D choice-encoding subspace (red arrow) captures all information about the decision. Note, for example, that the neuron 2 firing rate axis is nearly orthogonal to the choice axis, meaning that neuron 2 carries almost no information about choice. **(c)**. Projections onto the stimulus and choice subspaces reveal the time-course of information about stimulus and choice, respectively. These timecourses can be seen as temporal basis functions for the single-neuron PSTHs shown in (**a**). mTDR aims to recover these encoding subspaces even in the presence of additional components that take neural activity outside the plane spanned by these two axes, and is not restricted to 1D subspaces.

**Figure 2:**
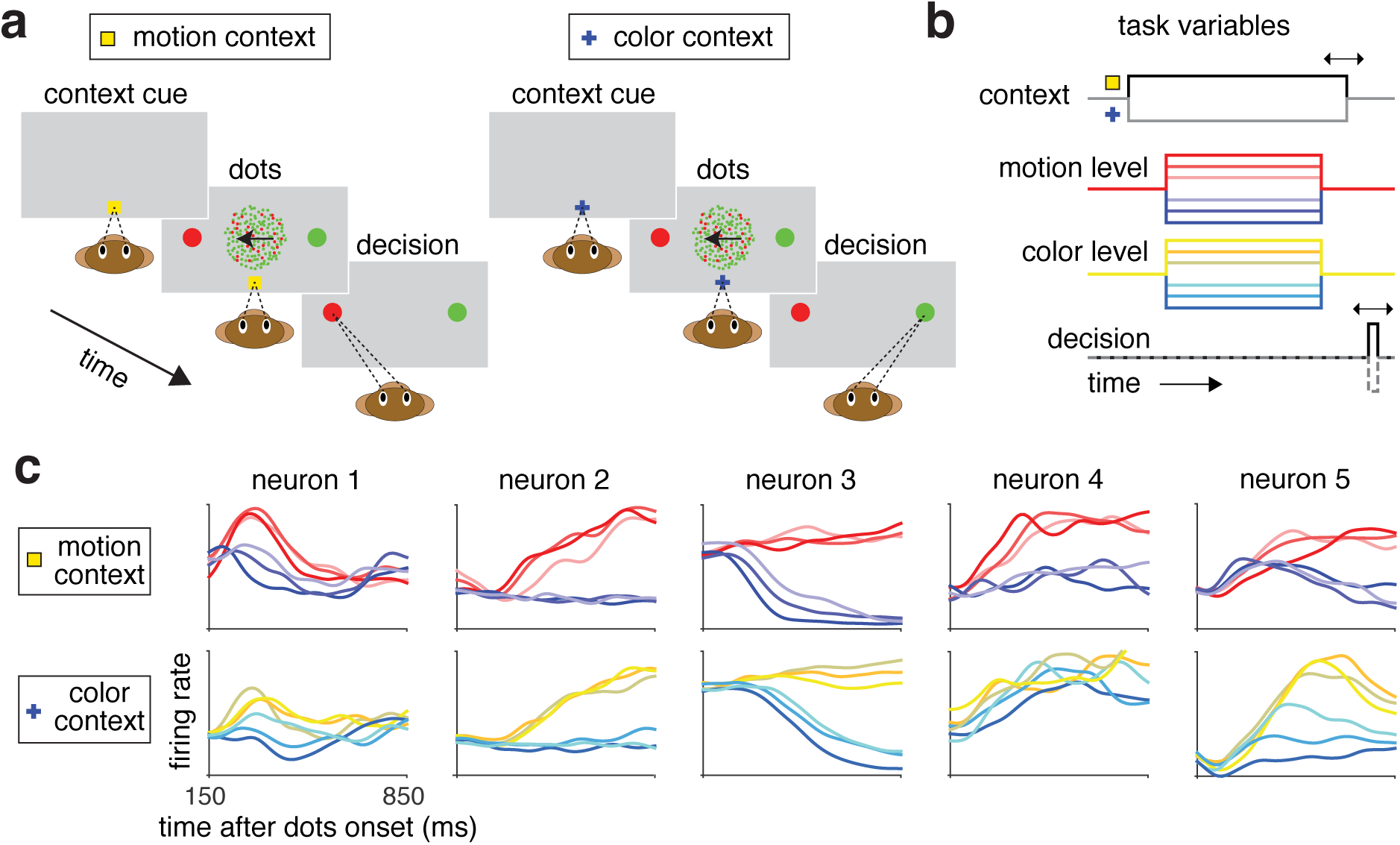
Context-dependent decision-making task and neural responses. **a**) On each trial, the animal was presented with a context cue (yellow dot or blue cross) indicating which dimension of the stimulus the animal is to attend to, followed by a stimulus of colored, moving dots. On motion context trials the animal is cued to respond to the dominant dot motion direction. In color context trials the animal is cued to respond to the dominant color of the dots. **b**) The strength of both the color (red / green) and motion (left / right) stimulus was displayed with one of six possible degrees of coherence, making for many possible task contingencies (2 choices × 2 contexts × 6 motion strengths × 6 color strengths = 144 possible combinations). **c**) PSTHs of representative neurons for monkey A. Motion context PSTHs were sorted by motion coherence and averaged over color coherence. Color context PSTH’s were sorted by color coherence and averaged over motion coherence. Red–indigo color scale indicates motion coherence where red indicates the preferred motion direction. Gold–blue color scale indicates color coherence where gold indicates the preferred color direction. Bolder colors indicate stronger coherence.

We also introduce a new unsupervised method, sequential principal components analysis, for decomposing multidimensional representations of task information into a sequence of axes that reflect the order in which information about each variable becomes available. This method reveals that multidimensional trajectories can be decomposed into an early phase with linear dynamics, followed by a later phase with rotational dynamics. We used model-based decoding under the mTDR framework to show that the transition between these phases corresponded to a sustained plateau in decoding accuracy for sensory as well as decision information, suggesting that the population did not continue to accumulate sensory information during the rotational phase.

Taken together, these results substantially extend the prevailing picture of decision encoding in PFC: rather than integrating evidence along a single dimension of population activity, with amplitude that reflects accumulated evidence ^17^, neural population activity enters a phase of rotational dynamics that maintains information about the choice as well as relevant and irrelevant sensory information over the entire course of a single trial. ^18–20^.

## Results

### Model-based targeted dimensionality reduction

To characterize population-level representations of information in PFC, we introduce a new method, *model-based targeted dimensionality reduction* (mTDR), which seeks to identify a set of dimensions of population activity that carry information about distinct task variables. We illustrate the basic intuition for mTDR with a hypothetical 3-neuron population in a perceptual decision-making task (Fig. 1). For this example, there are two task variables of interest: a sensory stimulus *x*_s_ and a binary decision variable *x*_c_. These variables modulate the firing rates in different ways and the modulations are time-dependent, producing a diverse pattern of population responses across conditions (Fig. 1a).

The population-level response can be examined in a 3-dimensional state space, where the coordinates of each axis correspond to the firing rates of each of the neurons (Fig. 1b). Although the full space is 3-dimensional, the trajectories traced out by these particular firing rates exhibit low-dimensional structure that is not apparent from the PSTHs alone (Fig. 1a). Specifically, the population activity is confined to a 2D plane defined by a pair of one-dimensional axes: a 1D “stimulus axis” (green arrow) captures information about the stimulus strength, while a 1D “decision axis” (red arrow) captures information about the choice. These axes capture all information about the task variables in the population. Projecting the population response onto each of these axes reveals a timecourse of information about stimulus level and choice, respectively (Fig. 1c).

The goal of mTDR is to identify these encoding subspaces from high-dimensional neural population data. For our three-neuron example, the mTDR model describes the time evolution of the population response **y**(*t*), a vector of 3 neural firing rates, as:

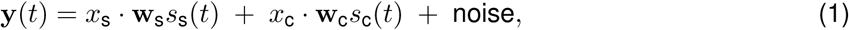

where *x*_s_ is the stimulus variable, which takes one of six values from [−3, −2, −1, +1, +2, +3] indicating the level of positive or negative sensory evidence, and *x*_c_ denotes the decision variable, which takes on values of *±*1, indicating a positive or negative choice. The activity vectors **w**_s_ and **w**_c_ are patterns of activity across the three neurons specifying the stimulus and choice axes (green and red arrows in Fig. 1b), and the time-varying functions *s*_s_(*t*) and *s*_c_(*t*) are temporal profiles for the activity along stimulus and choice axes, respectively (Fig. 1c).

Although the choice and decision subspaces in this example are both 1-dimensional, the mTDR model easily extends to higher dimensionality with an arbitrary number of task variables. Let **Y** denote a *neurons* × *time* matrix of firing rates for a single condition defined by task variables {*x*^(1)^, …*x*^(*P*)^}.

The mTDR model decomposes population activity as:

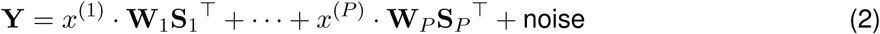

where **W**_*p*_ is *neurons* × *r*_*p*_ matrix whose columns span a *r*_*p*_-dimensional encoding subspace for task variable *x*^(*p*)^, and **S**_*p*_ is a *time* ×*r*_*p*_ matrix of temporal profiles that describe the timecourse of population activity within this subspace. (See supplementary Figure S1 for a schematic illustration of this model.) This model-based formalism represents a generalization of targeted dimensionality reduction ^1^, which allows us to identify both the number of activity patterns used to encode different variables and the timecourses with which these patterns are recruited. (For details see Methods and Supplementary Note S1).

### Population coding of task variables in PFC

To investigate population-level coding in PFC, we applied mTDR to neural data recorded from an area in and around the frontal eye fields (FEF) of two monkeys performing a context-dependent decision-making task ^1^ (see Methods, Experimental details) In this task, monkeys were presented with a visual stimulus that contained colored, moving dots on each trial (Fig. 2a). A context cue (yellow square or blue cross) appeared before each trial and instructed the monkeys to attend either to the color (red vs. green) or the motion (left vs. right) of the dots. In the color context, the animal had to attend to color and ignore motion, making a left (right) saccade if a majority of the dots were red (green). In the motion context, the animal had to attend to motion and ignore color, making a left (right) saccade if the dot motion was left (right).

Task difficulty was controlled by varying the fraction of red vs. green dots across 6 levels of color coherence (from “strong red” to “strong green”), and varying the fraction of coherently moving dots across 6 levels of signed motion coherence (from “strong left” to “strong right” motion), resulting in 6 × 6 = 36 unique stimulus conditions (Fig. 2b). The stimulus was followed by a randomized delay, after which the monkey was cued to indicate its decision by making an eye movement to one of the two saccade targets. Taking into account the 2 possible contexts and 2 possible decisions on each trial, there were 2 × 2 × 36 = 144 unique task conditions in total.

The classical approach to analyzing data from such experiments involves computing the mean firing rate, or peristimulus time histogram (PSTH), from subsets of the data, such as “all trials with the strong rightward motion and a rightward choice”. We will refer to these condition-averaged responses as *conditional PSTHs*. For this dataset, the conditional PSTHs of individual neurons exhibited heterogeneous tuning to the different task variables ^1^ (Fig. 2c). This hetereogeneity, and the fact that each neuron encodes a wide variety of task variables, makes it difficult to obtain a clear picture of the population-level representation of task variables from an examination of single-neuron PSTHs.

To overcome these limitations, we used mTDR to determine the dimensionality of population-level representations of the different task variables. The mTDR model included a regressor for each of 6 task variables: color strength, motion strength, context, and choice, as well as two additional terms for the absolute values color and motion strength. Absolute value terms were included due to the observations that some neurons displayed nonlinear encoding of stimuli, consistent with observations of nonlinear mixed selectivity ^13^. The model also included a term for the condition-independent time-varying firing rate, which reflects temporal modulation not due to the task variables (see Methods for details). To determine the dimensionality of the encoding of each task variable, we used a greedy selection method based on the Akaike information criterion ^21^ (AIC) that added dimensions based on their contribution to the model prediction performance. We validated this approach with simulation experiments and by using cross validation on the real data, which we found to slightly underestimate dimensionality due to the need to divide data into training and test sets (Fig. 3c; details in Supplemental Note S3).

**Figure 3:**
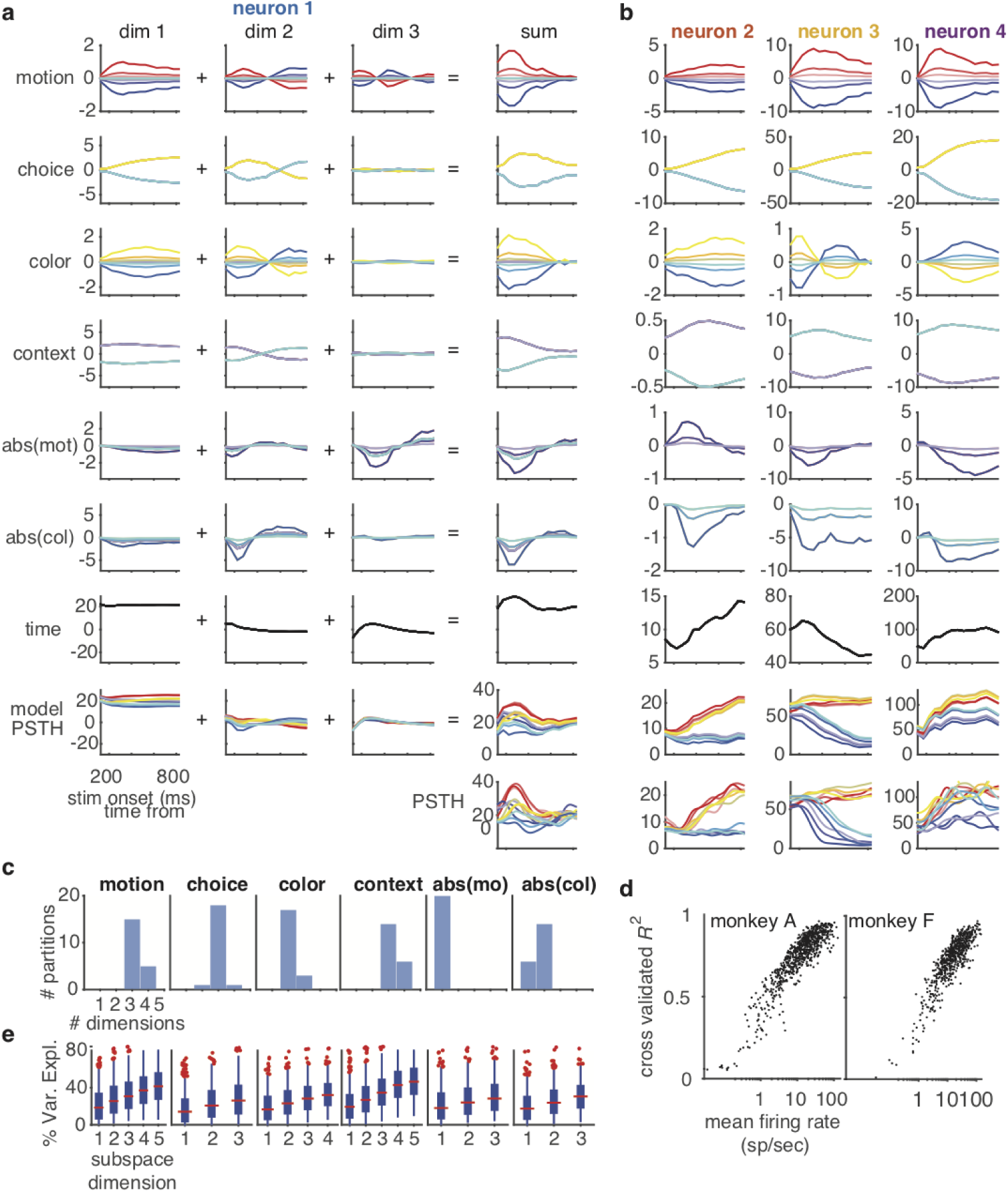
Model fit for monkey A. **a**) Example of a neuron’s fitted responses composed of a set of weighted basis functions (same as neuron 1 from Fig. 2c). These basis functions are shared by the whole population but are weighted differently for each neuron. Weighted basis functions are summed to form the neuron’s response to each task variable. The responses for each task variable are then added together to give the model reconstructed PSTHs (model PSTH). The conditional PSTHs of this neuron are shown for comparison. **b**) Summed responses for three additional example neurons (same as neurons 2–4, from Fig. 2c) which display a diversity of dynamics. **c**) Dimension estimation based on 5x 4-fold (20 estimates) cross validation. Dimensionality is slightly smaller than estimated using all data but is tightly distributed around a single estimate. **d**) *R*^2^ of the model reconstructions for the PSTHs as a function of mean firing rate for each neuron. **e**) Percent variance explained for PSTHs of each neuron by projection onto each subspace dimension. Red horizontal bars indicate the median. Box edges indicate 25th and 75th percentiles. Whiskers indicate positions of furthest points from median not considered outliers. Red dots indicate outliers with respect to a normal distribution. Dots have been horizontally jittered to aid with visualization. Colors in title text for (**a**) and (**b**) correspond to colors of markers in Figure 5.

We found that population-level representations of all task variables were at least two-dimensional, and at least three-dimensional in monkey A (Fig. 3; Supplemental Table 1; Supplemental Fig. S3). Figure 3a shows the variable-specific components revealed by mTDR for an example neuron. The first three columns show the timecourse of this neuron’s activity within the first three dimensions of the corresponding variable’s encoding space. The timecourses represent the columns of the temporal component matrices **S**_*p*_, scaled by the levels of each of the task variables *x*_*p*_ (eq. 2). Thus, each trace represents the inferred contribution of each dimension to the neuron’s PSTH from the different settings of the associated task variable. The rightmost column of Figure 3a shows the model-based estimate of the neuron’s net time-varying response to each task variable. Summing these responses together gives the model-based reconstruction of the neuron’s PSTH (“model PSTH”) for each task condition; this matches the neuron’s true PSTH to high accuracy (bottom). Because each neuron weights each dimension independently, the fitted model collectively accounts for a wide variety of conditional PSTHs (Fig. 3b). Note that the data were not temporally smoothed in pre-processing and no smoothness constraints were included in the model, indicating that the smoothness of the timecourses is a property of the data.

**Table 1:** Summary of estimated dimensionality of each task variable subspace.

| animal | motion | color | choice | context | abs(motion) | abs(color) |
| --- | --- | --- | --- | --- | --- | --- |
| monkeyA | 5 | 4 | 3 | 5 | 3 | 3 |
| monkey F | 5 | 2 | 3 | 4 | 3 | 2 |

To examine whether the model with estimated dimensionality provided sufficient richness to describe the diversity of PSTHs from the whole population, we calculated the *R*^2^ of our model for each neuron using held-out data. We found that the *R*^2^ of PSTH reconstructions increased with firing rate with the highest-rate neurons achieving *R*^2^ greater than .9 (Fig. 3d). The dependence of *R*^2^ on firing rate likely reflects higher signal-to-noise ratio in higher firing-rate neurons.

We also measured how much of the variance of the PSTHs formed from held-out trials could be explained by each of the learned subspaces alone (Fig. 3e). We defined each subspace by a set of orthonormal vectors ordered by the amount of variance explained (for details see Methods and Supplementary Note S4.1). We found that all dimensions contributed to the variance of at least some neurons, but that different neurons had their variance distributed differently across components (Fig. 5a,b). For example, for the decomposition of the activity of the neuron displayed in Figure 3a, dimension 3 of the abs(motion) axis has a higher loading than dimension 1 despite the fact that the first dimension describes most of the variance across the population.

**Figure 4:**
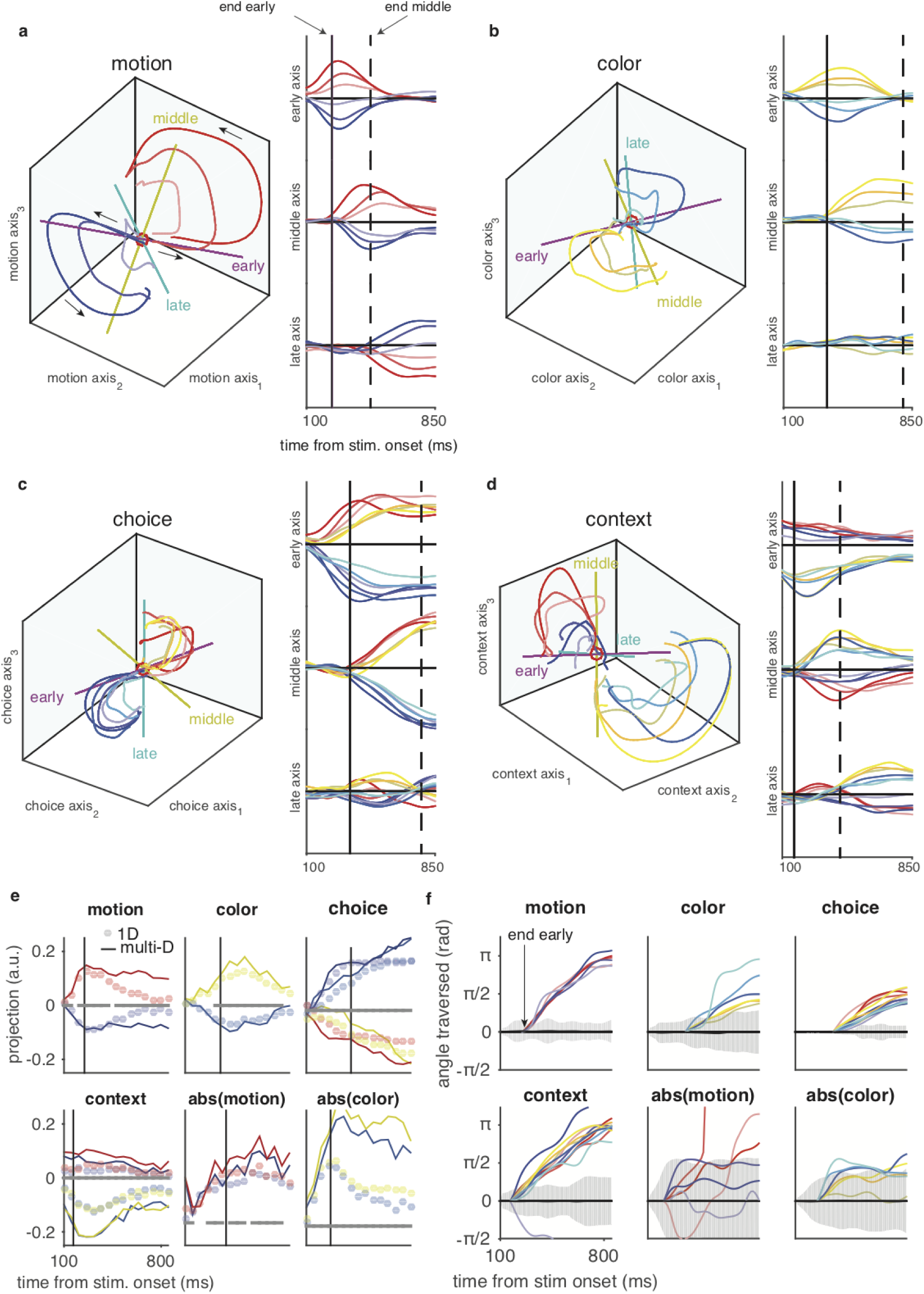
Projections of population PSTH’s onto latent encoding subspaces. Projections onto the first, second, and third principle-axes of the **(a)** motion, **(b)** color, **(c)** choice, and **(d)** context subspaces. Motion, color, and context subspaces have been orthogonalized with respect to the first dimension of the choice subspace. The choice subspace has been orthogonalized with respect to the context subspace. The context subspace has also been orthogonalized with respect to the motion and color subspaces. Details of orthogonalization are presented in Supplementary note S4.2. Color conventions are the same as those described in Figure 2. Red dots indicate the origin. Projected PSTH’s made from held-out data not used during parameter estimation. **a**) Projections of PSTHs onto the motion subspace, sorted by motion coherence and averaged over color coherence for trials where the motion stimulus was the active context. **b**) Projections onto the color subspace sorted by color coherence and averaged over motion coherence for trials where the color stimulus was the active context. **c**) Projections onto the choice subspace. Motion context trials are displayed with the same sorting and color conventions as displayed in (**a**). Color context trials are displayed with the same sorting and color conventions as displayed in (**b**). Only correct trials are displayed. **d**) Projections onto the context subspace using the same conventions as displayed in (**c**). Only correct trials are displayed. Colored axes in 3D plots indicate seqPCA axes. Solid vertical lines accompanying time traces indicate the time points where middle-axis variance starts to increase. Dashed vertical lines indicate the time points where late-axis variance starts to increase. Units of the ordinate are arbitrary but all time-trace axes are on the same scale. PSTHs were generated with ≈ 13 ms time bins and smoothed with a Gaussian window with standard deviation of ≈ 50 ms. **e**) Median encoding strength of pseudotrials onto the first three encoding axes of mTDR compared with the 1D subspace estimated by the max-norm method used by Mante et al. ^1^ (see Supplementary note S10 for details). For clarity, only trials with the strongest stimulus strengths are shown. Grey bars at *y* = 0 indicate time points where the mTDR projections had significantly stronger encoding across all stimulus levels than the 1D projections (left-tailed Wilcoxon signed-rank test, pFDR ^28^ controlled at .01). Multi-dimensional mTDR projections are larger than 1D projections at nearly all times for all task variables. **f**) Rotation angle traversed through rotational projection using jPCA. Angle was calculated starting from time when the projection transitions between the early and middle epochs. Coherent traversal across stimulus strengths that is consistent and monotonically increasing is an indication of rotation. Shaded areas are 95% confidence regions calculated using a maximum entropy method ^29^ under the null hypothesis of no population structure other than the empirical means and covariances across time, neurons, and task conditions.

**Figure 5:**
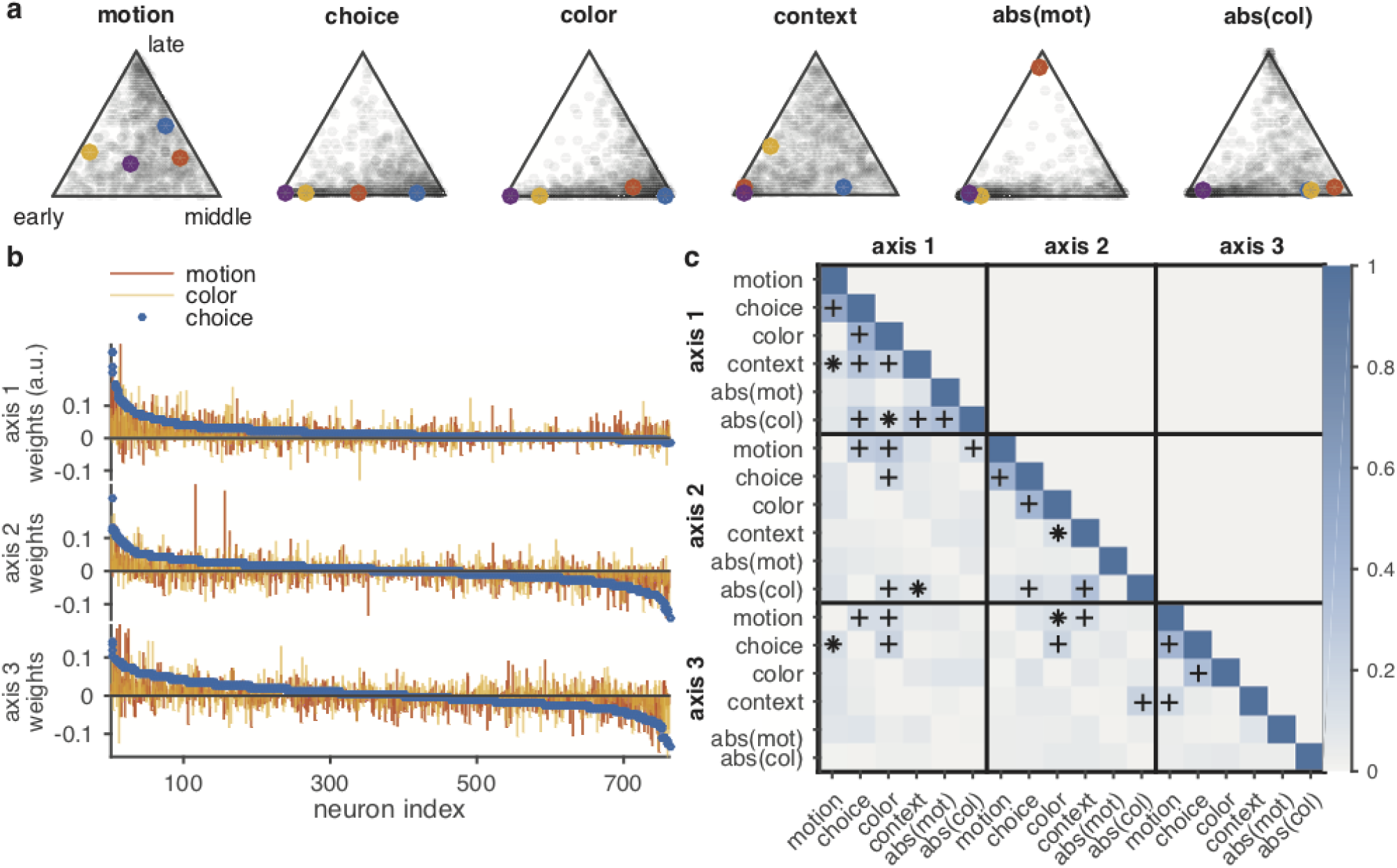
Distribution of variance within and between subspaces. (**a**) Proportion of variance among seqPCA axes. Each marker corresponds to one neuron. The position of each neuron indicates the distribution of variance from PSTHs across corresponding early, middle, and late axes. e.g. a point that lies closer to the “early” vertex of the motion plot has more of its motion-specific variance explained by the early axis while a point in the middle of the simplex has variance equally distributed across all axes. Darker regions indicate higher density of points. Colored dots correspond to cells displayed in Figure 3. (**b**)Weights of the top (in terms of variance explained) 3 axes for all cells for motion, color, and choice subspaces. Cell indexes are sorted according to the choice weights from most positive to most negative. (**c**) Magnitude of the Pearson correlation between top 3 subspace axes. The magnitude is used because the axes are only identifiable up to a sign. Markers indicate significant correlations controlled by the positive false discovery rate ^28^)(* *Q <* .01, +*Q <* .01). Null distribution is based on the positive half-Gaussian with zero-mean and standard deviation *σ*_0_ = 1*/n*, where *n* is the number of neurons. Significant correlations are most consistent between color-choice and motion-choice pairs.

These findings verify that the subspaces defined by the mTDR model capture high-variance dimensions and that the model describes a large fraction of the variance of the PSTHs for most neurons, despite the population representation being relatively low dimensional.

### State-space trajectories reveal dynamic encoding

To examine the dynamics of population-level encoding during this task we used projections of PSTHs from held-out data onto the estimated subspaces for motion, color, choice, and context (Fig. 4a–d; also see Supplementary movies; projections for abs(motion) and abs(color) are presented in Supplementary Fig. S9; monkey F projections shown in Supplementary Fig. S11; for details see Methods). Since all of the task variables for monkey A were estimated to have a dimensionality of 3 or higher, we will restrict our description of subspace trajectories to projections onto the three most significant axes.

Consistent with the findings in Mante et al. ^1^ using targeted dimensionality reduction (TDR), the dynamics in both the motion and color subspaces were qualitatively similar (Fig. 4a,b). However, in contrast to those findings, and the findings of others using a related TDR method ^8^, the encoding of information about the stimulus variables was not transient but persisted throughout the recording epoch, albeit along a changing set of dimensions at each point in time.

In order to examine when and how the stimulus encodings changed over time we developed a method for identifying an ordered set of axes that account for the variance of the projections sequentially in time. We term the method “sequential principle components analysis” (seqPCA) (see Methods, Supplementary Note S9). Using seqPCA we obtained 3 orthonormal axes that correspond to “early,” “middle,” and “late” epochs of the projections’ trajectories (labeled axes in left panels of Fig. 4a–d). The early axis accounted for the majority of the variance shortly after stimulus onset. Variability that is not described by the early axis but nevertheless emerges sometime after stimulus onset is captured by the middle axis. The late axis accounts for activity that is not accounted for by the early and middle axes, but is present as the epoch transitions from the stimulus presentation to the delay period. The late axis therefore may not exhibit perfectly sequential activation relative to the middle axis. Projections onto the seqPCA axes show clear times at which task variable information becomes available onto each axis (right-side panels in Fig. 4a–d). For all subspaces, we found that the early epoch is characterized by loading of the projections almost exclusively onto a single axis. In contrast, the middle and late epochs were distinctly two dimensional, or higher.

We found that the transience of the early axes in the stimulus subspaces resembles that of the stimulus encodings presented by the TDR method ^1^. Indeed, we found that our early axis was well correlated with the TDR axes (see Supplementary Note S10). It is therefore apparent that the existence of the middle and late seqPCA axes permit the stimulus information to persist throughout the stimulus viewing epoch. To show this, we compared projections onto the learned subspaces of the mTDR method with the 1D axes of the TDR method (see Supplementary Note S10). We found that while the loading of stimulus information appeared transient for TDR, the mTDR projections were both larger and more persistent at nearly all times during stimulus viewing (Fig. 4e, S12).

The encodings for motion and color for monkey A, and the motion encoding for monkey F, exhibit remarkable similarity (Fig. 4a,b and Supplementary Figs. S10a,b, S11a, S14a). Specifically, along the “early” axes stimulus encodings peak at around 300 ms after stimulus onset (Fig. 4a,b), peaking slightly earlier for motion than for color. Stimulus encodings begin loading onto the middle axes just prior to the choice trajectories (Fig. 4c). In all cases, the magnitude of the projections onto the seqPCA axes scales with the stimulus strength (Fig. 4b,d) and appear to statically encode the stimulus near the end of the stimulus presentation in a way consistent with delay-period encoding in parametric working memory seen elsewhere ^22–27^.

The population-level representation of choice, context, and other variables also exhibited multi-dimensional structure (Fig. 4c,d; Supplementary Fig. S9). We describe this structure and discuss its consequences in subsequent sections.

### Trajectories exhibit rotational dynamics

The projections for motion, color, choice, and context exhibit rotations after a short period of loading onto the early axes (Fig. 4a–d, left panels). This observation is supported by the fact that the trajectories are ≥ 2 dimensional during this period (Fig. 4a–d, right panels). While rotations are inherently ≥2-dimensional, the fact that we found trajectories to be ≥2-dimensional need not imply rotations. We therefore identified the plane of greatest rotation of the trajectories using jPCA ^18^ (Supplemental Fig. S10, Fig. S14), and observed clear rotational structure. The two dimensions of the jPCA plane accounted for a relatively large amount of the variance for all task variables (Fig. S15). Condition-shuffled projections yielded no apparent sequential or rotational structure (Supplementary Note S10, Fig. S16,Fig. S17).

In order to rigorously examine the presence of rotational dynamics, we examined the angle of rotation that the trajectories traversed from the beginning of the middle epoch to the end of stimulus viewing (Fig. 4f). We reasoned that for trajectories to be consistent with rotational dynamics they would have to have monotonically changing angle of rotation. We compared the angle of rotation to samples from the null distribution corresponding to the maximum entropy distribution with the same second order moments as the data ^29^ (Fig. 4f, see Supplementary note S5 for details). We found evidence for rotational dynamics in motion, color, choice, and context subspaces, although rotations were less consistent with the trajectories of the color encoding for monkey F (Fig.S11, Fig. S14, Fig. S18). These results indicate that rotational dynamics are not trivially present in these data and that we observe them in most of the linear subspaces examined.

Projections onto the subspaces for the absolute values of motion and color (abs(motion), abs(color)) were qualitatively different from those of the linear regression terms (Supplementary Fig. S9). While they clearly encoded the absolute values of the stimuli, evidence for rotational dynamics was not significant (Fig. 4f, Supplementary Fig.S10, Fig. S9).

### Neurons exhibit time-dependent tuning with stimulus encoding correlated with decision encoding

We used the mTDR model and seqPCA to examine the tuning properties of these cells. The encoding subspaces found by mTDR for motion, color, and choice, appeared to be correlated with one another (Fig. 5a). More specifically, the weights defining the motion and color bases were correlated with the choice weights but not with one another (Fig. 5c), indicating that motion and color representations both contributed to the choice encoding but that there was little interference between representation of motion coherence and the representation of color coherence.

Individual neurons exhibited complex mixtures of early, middle and late responses (Fig. 5b). While the population tuning of some task variables (abs(motion), abs(color)) were dominated by the early response none of the task variables were found to display clustering, but a continuous distribution of tuning across all three seqPCA axes. Late axes tended to explain less of the population variance, especially for color, choice, and abs(motion), but were responsible for explaining the majority of the variance for at least some neurons.

Also notable is the low density of cells near the early/late axis (i.e. left-arm of the ternary plots in Fig. 5b). This indicates that there are few cells that encode a task variable at the beginning and end of stimulus viewing but lose sensitivity to a task variable in the middle stimulus viewing. This implies that individual cells encode each task variable in continuous epochs, even if only transiently.

### Accurate stimulus decoding corresponds to transition in dynamics

Our generative model framework provides a natural setting for the decoding of population responses by maximum likelihood (see Supplementary Note S6). This allows our decoding analysis to be consistent with the results of dimensionality reduction. We can therefore investigate how and when the features of the low-dimensional trajectories translate into putatively perceived stimuli and behavior, and whether or not these features may be read out by downstream populations. We note that while decoding of task variables does not imply a causal role for the encoded variables in FEF function, decoding analysis does provide a clearer picture of the dynamics and fidelity of task variable encoding.

For decoding experiments with monkey A, we used a 4-fold cross validation in which we used 75% of the data to estimate parameters of the model and used the remaining 25% to produce 100 pseudosamples (with replacement) for decoding (for monkey F we used 2-fold cross validation with similar results). The resulting decoded values were averaged over pseudosamples and cross validation folds.

Stimuli could be accurately decoded within ≈150ms of stimulus onset for the motion stimulus and within ≈200ms for the color stimulus, roughly corresponding to the time of transition between the early and middle seqPCA axes (Fig. 6a). The decoded value of the stimuli were constant by the start of the middle epoch for both contexts and the variance of the decoding decreased dramatically up to this time (Fig. 6b). Thus, the change in population dynamics (early-to-middle transition) within the stimulus subspace was consistent with decoding accuracy and stability. The decoded values are slightly biased toward zero in the irrelevant context, suggesting some gating of information across contexts. Moreover, we found that the decoded stimulus values of the same sign were more closely spaced than the true stimuli. This effect is in keeping with the findings of Hanks et al. ^6^ where they showed that the encoding of stimulus evidence in FOF (a rodent analogue for the FEF) encodes accumulator values nonlinearly, with wider-than-linear spacing for moderate stimulus strengths.

**Figure 6:**
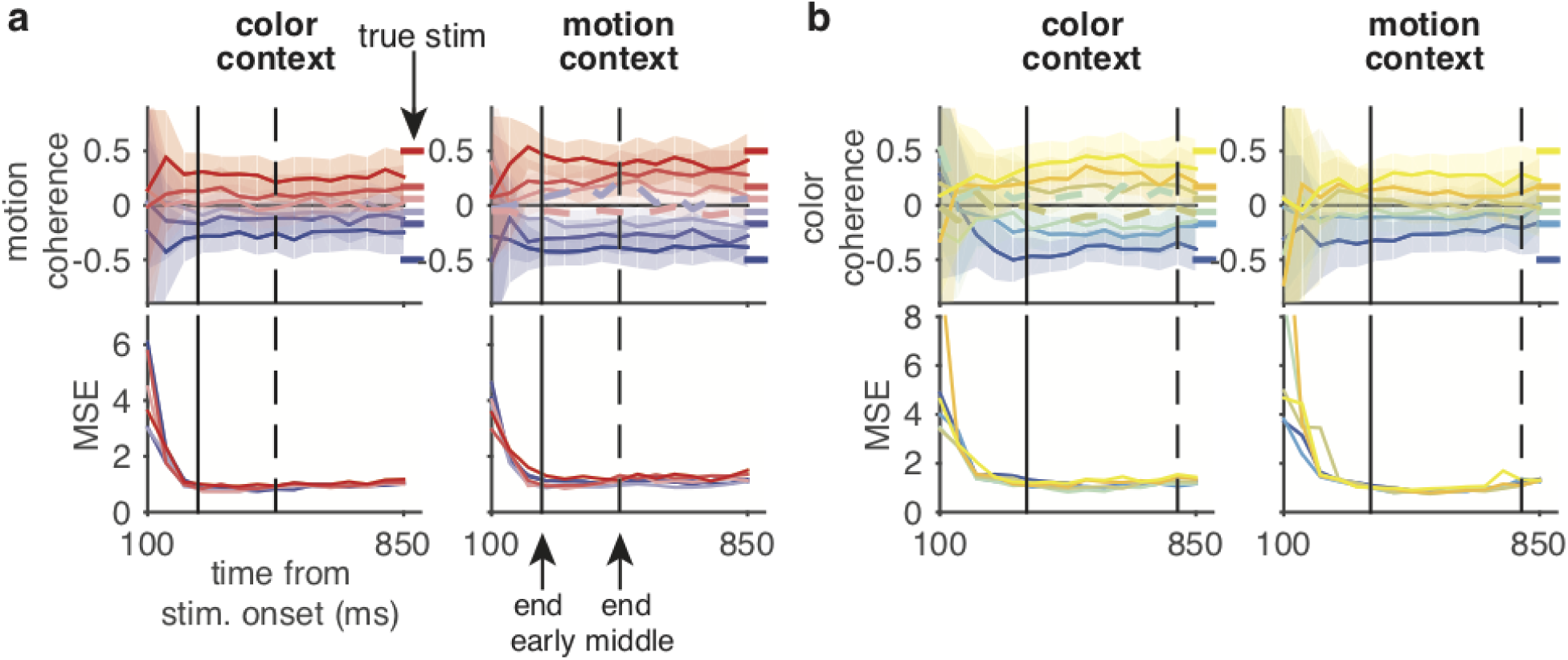
Instantaneous decoding of stimulus for monkey A. **a**) Top: Decoded motion coherence by mTDR model in both contexts. Bottom: Mean squared error (MSE) over time of motion coherence decoding across stimulus levels and context. MSE decreases precipitously, and then stabilize around the time of the first transition. **b**) Same as **a**) for color coherence decoding. Color conventions are the same as in Fig. 4. Shaded regions indicate 50% confidence intervals. Dashed lines indicate error trials from the corresponding context for the lowest stimulus strengths. 100 pseudotrials for each of 4-fold cross validation used for all analyses. Solid vertical lines indicate the time of early/middle axis transition for the corresponding stimulus subspace projections. Dashed vertical lines indicate the time of middle/late transition.

We also examined the decoded stimulus for error trials using the weakest stimulus strengths (dashed lines, Fig. 6c, Supplementary Fig. S19c). For these data only the weakest stimulus strengths had enough error trials to provide reliable statistical analysis ^1^. For monkey A, we found that the decoded stimulus values on error trials were similar to correct-trial decoding but were opposite in sign; suggesting that the origin of errors was (on average) an incorrect percept.

### Choice Decoding

We next studied how and when decision information became available in PFC and how the dynamics we observe in the encoding of choice translates into its decoding. In contrast to the decoding of stimuli, which are a continuous-valued variables, the choice variable was encoded in our model as a binary variable. Therefore, at each point in time we studied the log likelihood ratio (LLR) (for details, see Supplementary Note S6.3) of pseudotrials sampled from held-out data where the ratio was between the likelihood of a preferred, versus an anti-preferred, choice (Fig. 7a). Positive LLR indicates evidence in favor of a choice toward the preferred target and negative LLRs indicate evidence in favor of a choice toward the anti-preferred target.

**Figure 7:**
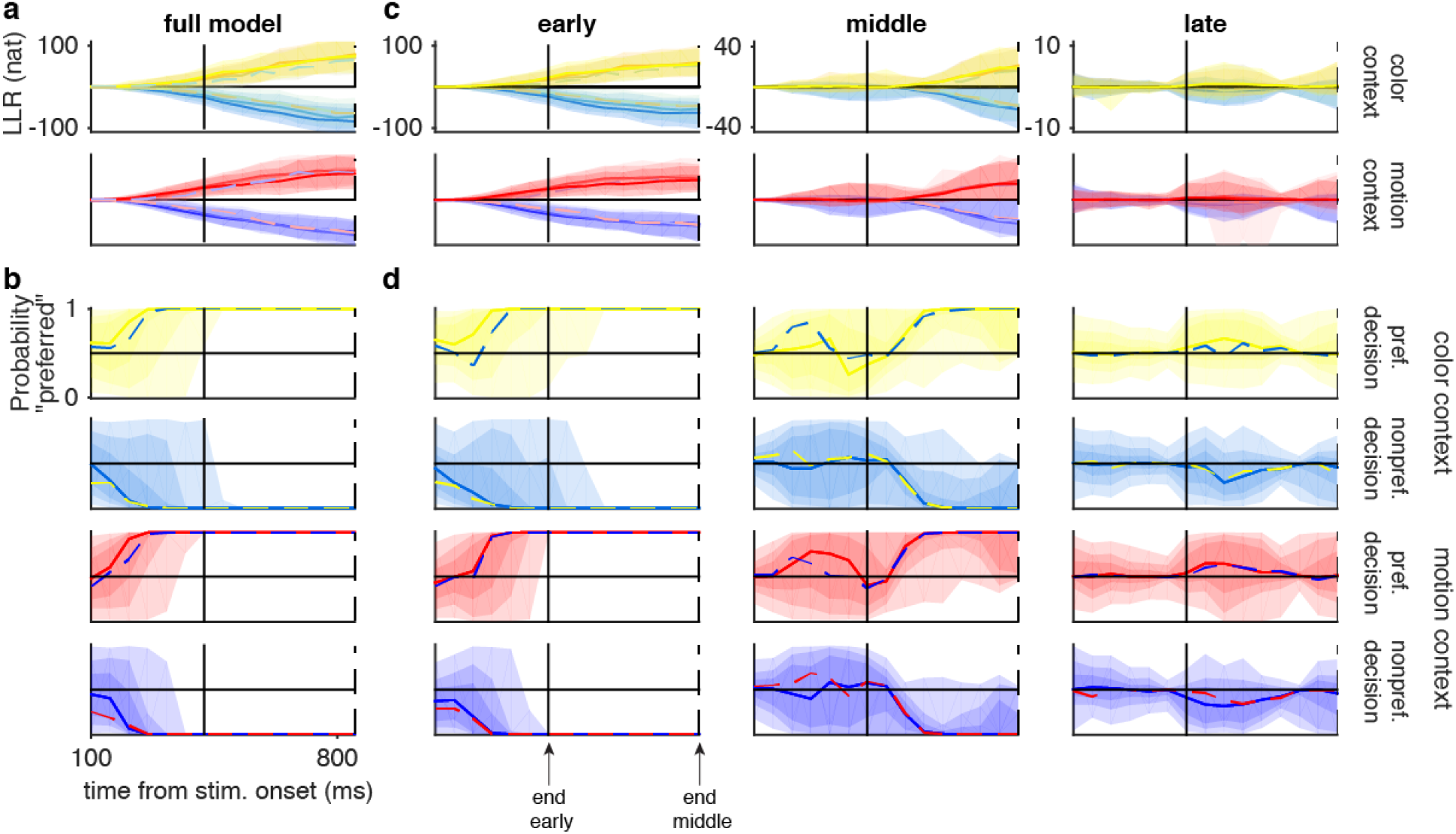
Instantaneous decoding of choice. **a**) Log-likelihood ratios (LLR’s) for monkey A in favor of a preferred choice using single pseudotrials from color - context (gold-blue, sorted by color coherence) and motion - context (red-violet, sorted by motion coherence) trials. Shaded regions indicate 95% quantile intervals for each stimulus strength. Solid lines indicate the median of correct trials. Dashed lines indicate median of error trials. **b**) Probability of a preferred choice based on corresponding LLRs combined over all stimulus strengths (see section S6.3 for details). Solid lines indicate median of correct trials. Dashed lines indicate median of error trials. Shaded regions indicate quantile coverage intervals of correct trials (light-to-dark: 95%,75%,50%). Color conventions are the same as in Figure 4. 100 pseudotrials for each of 4-fold cross validation folds used for all analyses. **c**) LLRs for in favor of a preferred choice where the choice subspace has been restricted to only the early, middle, or late axes. **d**) Probability of a preferred choice based on LLRs from (c).

The magnitude of the LLRs increased monotonically indicating an increasing strength of the decision signal over time. Although, the magnitude of the LLR did not differ strongly across context, direction of decision, stimulus strength, or whether the trials were correct or error trials (dashed lines, Fig. 7a). By transforming the LLRs into decision probabilities (Fig. 7b, see Supplementary Note S6.3) we were able to examine a moment-by-moment probability of the animal’s choice and ask when the decision is unequivocal and along which axes is the information available. We found that the choices could be discriminated with better than 95% accuracy as early as 300ms following stimulus onset for motion context trials and as early as 350 ms after stimulus onset for color context trials (Fig. 7b). This timing roughly corresponded to the time of transition between the early and middle seqPCA axes for choice. Similar results were observed for monkey F but timing was shifted slightly later (Supplementary Fig. S20). These results suggest that the animals had made their decisions on virtually every trial well before stimulus offset (at least 500ms before stimulus offset for monkey A and 100ms for monkey F) regardless of the stimulus coherence and that these decisions were coincident with a change in dynamics from a linear integration to a rotation within the choice subspace.

Interestingly, although the decoded choice for monkey F was somewhat more variable, a reliable decision signal was present from the first time point for many trials (Supplementary Fig. S20). This was particularly true for error trials, suggesting that either the decision was made earlier on average for error trials than for correct trials, or that the pseudosamples representing error trials are a mixture of both perceptual errors and “lapse” trials in which the animal did not attend the stimulus and made its choice by guessing.

We next examined the LLRs while restricting the choice subspace to only the early, middle, or late axes. The LLRs displayed the same invariance to choice, stimulus strength, context, and correct/error trial identity as the full model. For both monkeys, the early axis provides the majority of the available information about the decision and decoding along the early axis alone is nearly as accurate as when we use the full model (Fig. 7d and Supplementary Fig. S20d). However, the middle and late axes also displayed information about the choice later during stimulus viewing. Because we can decode the animals’ decisions with the early axis alone, it would seem as though the middle and late axis information is redundant and it is unclear what the purpose of these axes are. Similar multidimensional encoding of decision has been observed previously in premotor cortex ^30^.

### Context Decoding

We also examined the context signal using the same LLR method as our analysis of choice (Supplementary Fig. S21a, S22a). Similarly, the context evidence did not differ strongly across decision, stimulus strength, or whether the animal provided a correct or incorrect response. Transforming the LLRs into a probability of the perceived context (Supplementary Fig. S21b, S22b) showed that the correct context could be identified for both monkeys on the majority of pseudotrials from the first time point, which is consistent with the fact that the context cue was presented 650 ms before stimulus onset ^1^. These patterns all appear to hold for LLRs obtained with error pseudotrials as well as when decoding was restricted to only the early, middle, and late subspaces (Supplementary Fig. S21c,d, S22c,d). These findings demonstrate that accurate context information was available in PFC for the vast majority of both correct and error trials, suggesting that confusion about context was not a significant source of errors for either animal.

## Discussion

Our analyses have shown that PFC encodes individual task variables in distinct multidimensional sub-spaces within which the representation changes over time. The population activity patterns representing each task variable tended to follow a stereotyped pattern of 1D/linear encoding, followed by rotational dynamics. Our ability to make these observations was enabled by a new method of dimensionality reduction that is based on a generative model of the data.

We found that the dynamic nature of encodings in PFC requires multiple dimensions of neural population activity for accurate characterization. In particular, only multidimensional encoding, as opposed to 1D encoding, captures the persistence of stimulus information in PFC throughout the stimulus-viewing epoch (Fig. 4a,b). This finding complements the original report of these data ^1^, as well as results reported by others using similar 1D targeted dimensionality reductions methods ^8^. Our results suggest that previously reported transient stimulus encoding in PFC is only consistent with the early encoding axis (Fig. 4e). Our observations resemble multi-dimensional stimulus coding that mixes transient and persistent components ^30^ as well as population code “morphing” ^16^, where the optimal weights for decoding from population activity change over time, although the results shown here are on a time scale that is nearly an order of magnitude faster than previously reported.

While we validated our method for identifying the “true” dimensionality of the data using simulation experiments, it is unclear if the dimensionality would differ under different experimental conditions. Specifically, the dimensionalities we learned are likely to be influenced by a variety of factors ^31^ including the sample size, the fraction of neurons observed, the intrinsic model dynamics, and the task complexity ^32^. Some of these factors may explain the differences in dimensionality between the two animals in the present study, where the dimensionalities of monkey F were lower than monkey A in correspondence with smaller sample sizes and fewer recorded cells. However, we would like to emphasize that during the early encoding, nearly all trajectories are 1D and only afterward are ≥2D. This may be a direct reflection of the rotational nature of the trajectories following the first transition, where rotations are inherently ≥2D since they require both sine and cosine parts.

The mTDR method is distinct from unsupervised methods like PCA or factor analysis in that it uses information about the trial structure in order to perform dimensionality reduction. The method is also distinct from previously proposed supervised methods ^1;25–27;33^ in its use of an explicit statistical encoding model to describe the transformation from task variables to neural activity patterns. This distinction not only allows us to make predictions of population responses to experimental contingencies not observed in the data (something not possible for methods based on the conditional PSTHs like dPCA without model-based interpolations ^27^) but it allows us to apply the tools of probabilistic modeling and inference to estimate both the model parameters and the dimensionality of the encoding.

Our descriptive model of neuronal responses (eq. 5) is similar in principle to that used by Mante et al. ^1^ and other linear regression models used previously (see examples ^22;34–36^). However, ours is distinguished in its explicit specification of low-rank regression parameters and neuron-specific noise variance. Future iterations of our model may be improved by accounting for nonlinear mapping of stimuli onto neuronal responses ^22^, by modeling of noise correlations between simultaneously recorded neurons, and accounting for variable trial lengths.

Much theoretical development has rested on the notion that single-neuron spike rates map onto an evidence accumulator but recent evidence in the frontal orienting field (FOF, a rodent analogue of the FEF) has challenged this view ^6^, suggesting that this region can be better described as maintaining a running motor plan (saccade for FEF, orienting for FOF) based on the evidence accumulated so far ^6 7^. While our analysis does not aim to suggest a causal role of FEF, the results of the present study could be interpreted as supporting this view, where the early dynamics represent an evolving decision and the rotational dynamics indicated an evolving motor plan, but more work is needed to determine the precise role of FEF.

### Functional significance of sequential subspaces

Our analysis revealed temporally segregated dynamics with early-axis, linear activity transitioning to middle- and late-axes, with rotations dominating by around 200–400 ms after stimulus onset (Fig. 4, Fig.S10). The temporal separation of the linear and rotational subspaces suggest that these are sub-spaces within which distinct computations are evolving ^18;20;37^ or have independent sets of down-stream targets ^19^.

With the present data we can only speculate about what the nature of these different computations must be but the analysis presented here indicates the possibility that the early subspace is a correlate of the temporal window within which decision making is performed. For example, the time-frame of transition between early and middle epochs is consistent with the time frame within which we can decode the animals’ decisions from single pseudotrials (Fig. 7 and Supplementary Fig. S20). This time frame is consistent with the time-frame of saturation of the chronometric curve for the traditional moving dots task ^38–40^, is consistent with the distribution of step times in the stepping model of evidence accumulation ^41^, and is consistent with early weighting of evidence in visual discrimination tasks ^42^. This evidence suggests that the transition from linear to rotational dynamics is a correlate of decision commitment.

In premotor cortex, a similar sequence of dynamics has been observed in population activity that corresponds to distinct “preparatory” and “movement” epochs ^18–20;37^. However, the latter findings were isolated to motor and premotor areas while ours were from dorsolateral PFC, localized around FEF ^1^. In addition, in the premotor and motor cortex studies the transitions in dynamics could be linked directly to an overt action (arm movement) while our animals would not have made an overt action (saccade to target) until 300-1,500 ms after the end of our analysis epoch ^1^. Therefore, if the animal has made its decision then it would have done so only covertly.

These distinctions, however, may very well be superficial. The qualitative features of our results reflect observations in motor cortex strikingly well ^18–20;37^ suggesting that common mechanisms may be at work in both motor execution and decision making. Indeed, FEF is defined as a region that elicits eye movement under stimulation ^43;44^ and has been implicated as a region important for visual decision making ^1;4;6;9–11;11;12;14;15^, oculomotor planning ^45^, and covert visuospatial attention ^46;47^. Thus, although FEF is not a motor region *per se* we may think of FEF as itself a premotor area responsible for visuospatial attention and motor planning concomitant with decision making ^4;6;7;9^. While the dynamic transitions in our analysis could be interpreted as signaling decision commitment rather than an action plan ^6^, it seems reasonable to view the distinct spatiotemporal partitioning of dynamics we find in the present study as signaling a covert action preparation that reflects the upcoming saccade, in analogy with the spatiotemporal transitions observed between preparatory and movement periods seen in premotor cortex ^18;19;37^. Single-trial population analysis and analysis of trajectories that extend into the delay and saccade epochs of these experiments may shed light on how the dynamics we observe reflect the animals’ decisions.

Some subspaces lack a distinct late component (eg. color and choice subspaces for Monkey A, Fig. 4a,c). However, it is possible that the middle seqPC for some task variables is serving a similar role as the late seqPC for others; preparing the network for a new set of targets and storing the memory of the stimuli as persistent activity over the course of the delay period. Indeed, the number of seqPCâĂŹ s needed to describe the population activity may be a reflection of the rate at which rotations twist into new encoding directions and therefore reflect a quantitative difference in encoding rather than a qualitative one. Future work should be aimed at identifying the significance of the dimensionality of the encoding relative to changing dynamics.

Finally, the nature of dynamic encoding for the context variable remains mysterious. Context encoding for both animals displayed clear and consistent dynamics (Fig. 4d,S11d) including rotations (Fig. 4f). Furthermore, while most of the predictive capacity of the context encoding lies in the early subspace (Fig. S21, S22), where context is encoded throughout the stimulus viewing period, context encoding at the single-neuron level is broadly distributed across the early, middle, and late axes (Fig. 5b, S13), indicating that some neurons do not encode context until well after stimulus onset. Further work is needed to determine what, if any, function these dynamics serve in decision making and memory.

### Differences in encoding between animals

The two monkeys in this study displayed similar, but not identical, encoding properties. For example, the trajectories through the motion subspaces are strikingly similar (Fig. 4a, Supplementary Fig. S11a) but we found obvious differences between the encoding trajectories for color (Fig. 4b versus Fig. S11b). For monkey A the color trajectories closely resemble the trajectories for motion (Fig.4a,b) while for monkey F the color trajectories display no obvious rotational component (Supplementary Fig. S11b). Choice and context trajectories in monkey F appear to be similar to those of monkey A (Fig. 4e,g and Supplementary Fig. S11e,g) but display less pronounced rotations, (Fig, S10, Supplementary Fig S14). These across-animal differences verify that rotational dynamics are not trivially present in these data and while it is unclear precisely what function they serve they are a potentially important feature of encoding in PFC. The abs(motion) and abs(color) trajectories in both monkeys appear to follow similar dynamics, with an early response peaking ≈200-300 ms and a later response that appears to tonically encode the magnitude of the stimulus (Fig. S9a,c and Supplementary Fig. S11e,f).

These differences are reflected in the our decoding results as well. While the qualitative results of stimulus and choice decoding appear to hold across animals, motion decoding appears to be more precise for monkey F (Fig. S19) than for monkey A (Fig. 6) and the 1D color decoding in monkey F is far more sensitive to the animals’ choice and the quality of decoding is relatively poor (Fig. S19). We also found that the transition between early- and middle-epoch decoding accuracy is less dramatic for monkey F than for monkey A (Fig. S19).

While the reason for differing dynamics between the color encoding for monkey F and the other stimulus encodings is unclear we do have some behavioral clues as to its effect. For example, the color-context psychometric curve for monkey F was somewhat more shallow than for motion as well as for both motion or color for monkey A (Extended data Fig. 2d in ^1^), and motion served as more of a distraction during the color task for monkey F than for monkey A, suggesting that color discrimination task was more difficult for monkey F. Furthermore, we found that the decoding accuracy for color in monkey F was considerably worse than for monkey A (Fig. 6, Fig. S19) suggesting that color information was more poorly represented in PFC for monkey F. Although not definitive, together these results suggest that monkey F may have had more difficulty with color coherence perception and that the encoding dynamics we observe are a correlate of perceptual uncertainty. Future experiments could be aimed at examining this hypothesis.

### Decoding of error trials suggests sources of errors

There are three ways that the animals may commit an error: the animal perceived the wrong stimulus (e.g. perceived left motion on a right-motion trial); the animal was confused about the context (e.g., made its decision using the color information in the motion context); or the animal made a random choice, independent of context cue or stimulus (i.e., a “lapse” trial). The results of this analysis indicate that the most likely of these scenarios, for monkey A at the weakest stimulus strengths, is that the animal perceived the wrong stimulus. We showed that the decoded context on most trials was the correct context, suggesting that the correct context was also the perceived context (Fig. S21), ruling out confusion about which stimulus the animal was supposed to attend. We also showed that lapse errors do not contribute significantly to the animal’s behavior since the LLR in favor of the executed choice indicates evidence in favor of the choice made on error trials that rose as fast as the correct trials, and follows essentially the same time course (Fig. 7a,b), where lapse errors would be indicated by a LLR that signals a decision earlier than correct trials. Indeed, the psychophysical curves of monkey A suggest a small lapse rate, if any ^1^. Finally, the decoded stimulus on error trials indicates that the perceived relevant stimulus on error trials was of the opposite sign as the stimulus that was presented (Fig. 6c). Together, these observations indicate that the error trials are characterized by a deliberated decision based on an incorrect perception of the relevant stimulus. A more direct trial-by-trial analysis of simultaneously recorded neurons would be useful in probing this hypothesis.

The results for monkey F are more difficult to interpret. The decoded stimuli for error trials appear to be close to 0, indicating an ambiguous stimulus (Fig. S19b). Furthermore, the choice signal on error trials appears to be present earlier on average than on correct trials and is present on some trials as early as the first time point (Fig. S20b) suggesting that the animal may have made its decision before even viewing the stimulus, suggesting that a significant source of errors for monkey F are lapses.

Given the present data, it may be impossible to distinguish the neural correlates of the animals’ choices from neural correlates of motor planning for the eventual saccade. Recent work has shown that there may be independent cortical signals for evidence accumulation and decision commitment in other cortical areas ^42^. It may therefore be difficult using data of this kind to distinguish between a deliberate effort to make a stimulus discrimination and the formation of a motor plan ^34^.

Nevertheless, the results presented here demonstrate the utility of mTDR for the analysis of neuronal population data and provide a description of PFC dynamics that should serve as important constraints on future models of the mechanisms of PFC function.

## Methods

### Detailed description of model

#### High-dimensional description of observations

Our model describes trial-by-trial neuronal activity with a linear regression with respect to the task variables. We assume that the activity of the *i*^th^ neuron *y*_*i,k*_(*t*) at time *t* on trial *k* can be described by a linear combination of *P* task variables 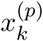, *p* = 1, …, *P* (eg. stimulus variables, behavioral outcomes, and nonlinear combinations thereof), such that

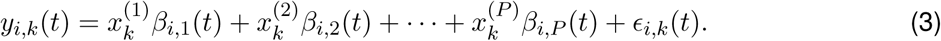

where the *P* values of the task variables 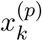 are known, the *β*_*i,p*_(*t*) are unknown coefficients, and *ϵ* _*i,k*_(*t*) is noise. This basic model structure is identical to that of the regression model used in ^1^ and has been successfully employed in characterizing neuronal activity of single neurons in other studies of perceptual decision making ^23 48^. In cases where we include a time-varying mean rate that is independent of the task variables, we define 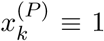 for all *k*, and the *P*^*th*^ component becomes the time-varying mean.

To represent all neurons simultaneously, we concatenate the responses into a vector **y**_*k*_(*t*) and write

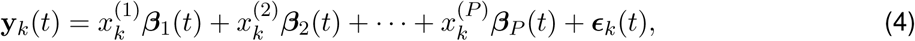

where **y**_*k*_(*t*) = (*y*_1,*k*_(*t*), …, *y*_*n,k*_(*t*))^┬^, ***β***_*p*_(*t*) = (*β*_1,*p*_(*t*), …, *β*_*n,p*_(*t*))^┬^, and ***ϵ***_*k*_(*t*) = (*ϵ*_1,*k*_(*t*), …, *ϵ*_*n,k*_(*t*))^┬^.

For trial epochs of duration *T* we can regard all observations on a given trial to be a matrix, **Y**_*k*_ = (**y**_*k*_(1), …, **y**_*k*_(*T*)), giving the observation model

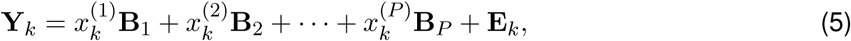

where **E**_*k*_ = (***ϵ***_*k*_(1), …, ***ϵ***_*k*_(*T*)), and **B**_*p*_ = (***β***_*p*_(1), …, ***β***_*p*_(*T*)). For the present study, we assume the noise is normally distributed ***ϵ***_*k*_(*t*) ∼ *𝒩* (0, **D**^−1^) for all trials *k* and times *t*, where **D** = diag(λ_1_, …, λ_*n*_) is a *n* × *n* diagonal matrix of noise precisions.

#### Low-dimensional description of observations

With no additional constraints our observation model (5) is extremely high dimensional and is effectively a separate linear regression for each neuron at every time point. This would only be a sensible model if we believed that neurons were not in fact coordinating activity between each other or across time. To define our low-dimensional model we can describe each **B**_*p*_ by a low-rank factorization, i.e. **B**_*p*_ = **W**_*p*_**S**_*p*_, where **W**_*p*_ and **S**_*p*_ are *n* × *r*_*p*_ and *r*_*p*_ × *T* respectively, where *r*_*p*_ = rank(**B**_*p*_). Equivalently, we can say that *r*_*p*_ is the dimensionality of the encoding of task variable *p*. This is equivalent to saying that the characteristic response of each neuron to the *p*^th^ task variable can be expressed as a linear combination of *r*_*p*_ weighted basis functions 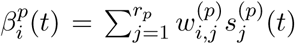, where *r*_*p*_ is the dimensionality of the encoding, 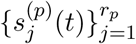 are a common set of time-varying basis functions, and 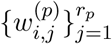 are neuron-dependent mixing weights.

### Marginal estimation of model parameters

The goal of inference is to estimate the factors of **B**_*p*_ and the ranks *r*_*p*_. Our proposed estimation strategy, for computational and statistical efficiency, is to estimate only one set of factors ({**W**_*p*_} or {**S**_*p*_}). This is possible when we integrate out one set of factors. For example, if we define a prior probability density over the mixing weights *p*(**W**), then for data likelihood *p*(**Y**|**W, S**) the marginal likelihood of the matrix of time-varying basis functions **S** can be obtained by

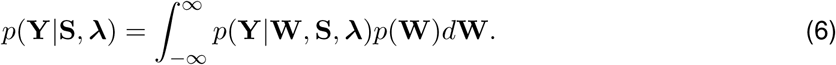

In principle, either set of factors may be selected for marginalization. In practice however the set of factors with lowest dimension should be selected to keep computational costs low. In this paper we focus on the case where *T* ≪ *n* and we therefore will estimate the set of weights {**S**_*p*_} while integrating over {**W**_*p*_}. The fact that either set of factors may be determined in this way means that there is a duality between rows and columns imposed by this model that is similar in principle to the duality between factors and latent states for probabilistic principle components analysis ^49^.

If we let the noise distribution and prior distribution of **W** both be Gaussian then we can use standard Gaussian identities to derive the marginal density *p*(**Y**|**S, λ**) and the corresponding posterior density *p*(**W**|**Y, S, λ**). A simple starting assumption would be to let all elements of **W** to be independent standard normal, (i.e., 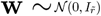 where 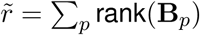). We therefore assume that the weights are a priori independent and that the noise variance is independent across both neurons and time. In principle, our framework supports the application of more structured priors and noise covariances, but we will not explore more elaborate models in this paper. Further details are developed in Supplemental Section S1

### Experimental details

A detailed description of these data have been published previously ^1^. Briefly, two adult male rhesus monkeys were trained to perform a context-dependent 2-alternative forced-choice visual discrimination task. At the beginning of each trial the monkeys were cued (Fig. 2a) to respond to either the motion or the color parts of the stimulus. After the context-cue presentation two targets appear for 350 ms, followed by 750 ms presentation of the stimulus. The stimulus was then followed by a randomized 300– 1500 ms delay after which the monkey was cued to indicate its decision with a saccade to either of the two targets.

Electrophysiological data were recorded from tungsten electrodes implanted in the arcuate sulcus in and around the frontal eye field (FEF). Electrodes were lowered two at a time into adjacent grid holes and were advanced until at least one single-unit could be isolated, although some trials yielded multiu-nit activity. All recorded units were included in the analysis. Spike sorting was conducted by clustering based on principle components analysis using the Plexon offline sorter (Plexon Inc., Dallas, TX). Each isolated cluster was functionally treated as a unit. Some clusters did not correspond to well discriminated, single-unit activity and were therefor deemed multi-unit activity.

All analyses presented in this paper used spike counts binned at 50ms (for model fitting and decoding) or 12.5 ms (for display of projections, jPCA, and PSTHs). All data were analyzed with custom scripts written in MATLAB (The MathWorks, Inc., Natick, MA).

### Model structure

Examination of the PSTHs revealed that stimulus encoding was asymmetric (eg. unit 2 in Fig. 2c), such that the encoding of the stimulus strength was stronger in one direction than the other. This suggested that the absolute value of the stimulus strengths should be jointly modeled with the linear encoding of the stimuli. Model fits using terms for the absolute value of the stimuli resulted in smaller AIC than model fits with only linear terms (Monkey A: AIC_linear_ = 9.79 × 10^7^, AIC_abs_ = 7.33 × 10^7^, Monkey F: AIC_linear_ = 8.065 × 10^7^, AIC_abs_ = 8.0628 × 10^7^).

### Cross validated variance explained

To asses the variance in the population responses that is explained by our method we conducted 4-fold cross-validation (CV) where, on each fold of CV, we used a randomly selected sample of 75% of the trials as training data to estimate the parameters of the model. Using the remaining 25% of the trials as test data, we made PSTHs for every possible task variable contingency for correct trials (total of 144 conditions). The reported variance explained was averaged over the four CV folds.

When assessing variance explained, the population PSTH’s for each condition was averaged over all extraneous task variables. For example, to assess the variance explained by the motion subspaces we averaged the PSTHs over all task variables except motion. We therefore had 6 sets of PSTHs for each neuron that were projected onto the motion subspace.

To determine if the variance that was explained by the estimated subspaces was greater than chance we compared the observed variance explained to the distribution of variance explained obtained by random projections. As a serrogate null distribution we generated 500 samples for each task variable of random projection weights from a normal distribution and calculated the explained variance for each sample. We then asked what the probability was of the observed explained variance being larger than the explained variance of the random projections for each neuron. We found that many neurons exceeded the 95% Bonferroni-corrected significance threshold across nearly all dimensions.

### Sequential PCA (seqPCA)

The seqPCA algorithm identifies an orthogonal basis on which variance of a *D*-dimensional trajectory is sequentially explained. The algorithm starts by calculating the variance explained by the first singular vector of a sequence of *D* × *t* data matrices **Y**_*t*_, where *t* indicates the number of time points included in the data. As the number of data points increases, the first singular vector explains a larger proportion of the variance,*p*_1,*t*_, until trajectories change direction, after which *p*_1,*t*_ decreases. The *t* at which *p*_1,*t*_ reaches its peak is considered a transition time and the left singular vector at this time is considered the first seqPC. Variability explained by this axis is subtracted from the data and the procedure is repeated to identify the 2nd seqPC, and so on. For details, see Supplementary Note S9.

The seqPCA algorithm displays some sensitivity to noise by making peaks in *p*_1,*t*_, difficult to identify. However, moderate smoothing (Gaussian window, 50ms width) of the trajectories appeared to mitigate this effect. Greater robustness may be offered by translation of this algorithm into an optimization framework ^50^. A related method has been developed for identification of sequential motifs of spike rasters ^51^.

## Acknowledgements

J.W.P. was supported by was supported by grants from the McKnight Foundation, Simons Collaboration on the Global Brain (SCGB AWD1004351) and the NSF CAREER Award (IIS-1150186). V.M. was supported by the Swiss National Science Foundation (SNSF Professorship PP00P3-157539), the Simons Foundation (to William T Newsome and Valerio Mante, Award 328189), the Swiss Primate Competence Center in Research, Howard Hughes Medical Institute (through William T Newsome, Investigator), and the DOD | USAF | AFMC | Air Force Research Laboratory (AFRL): William T Newsome, agreement number FA9550-07-1-0537

## Author Contributions

M.C.A and J.W.P. developed the model and performed data analysis. V.M. conceived and conducted the experiments and collected the data. All authors helped with the interpretation of data and writing of the paper.

## Data availability

Data are available from the corresponding author upon reasonable requests.

## Code availability

Demo code for the mTDR method is available for Matlab at http://www.mikioaoi.com/samplecode/RDRdemo.zip

## Supplementary Information

### S1 Derivation of model likelihood

The response of neuron *i* on trial *k* can be described by

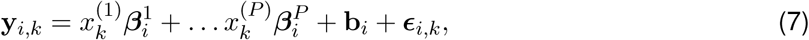

where all vectors are of length *T*, **b**_*i*_ is a constant vector representing a condition-independent mean, and ***ϵ***_*i,k*_ is noise. The low-dimensional description of the response is represented by a factorization of the vectors 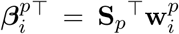 where, if *r*_*p*_ is the dimensionality of the subspace for task-variable *p* then 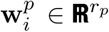 is a neuron-specific vector of weights and 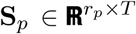 is a matrix of *r*_*p*_ time courses shared by all neurons. The basic model structure is graphically depicted in Fig. S1. If we let 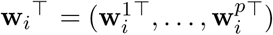, and **S** be a block-diagonal matrix

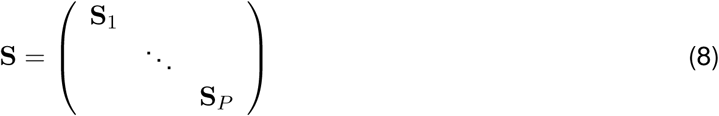

then we can rewrite equation (7) as

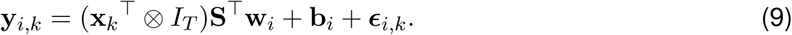

If **y**_*i,k*_ and **x**_*k*_ are the observed response and task variables on trial *k* then the collection of all observations for this neuron **y**_*i*_^┬^ = (**y**_*i*,1_^┬^, …, **y**_*i*_,*K*_*i*_ ^┬^) can be described in terms of all corresponding task variables 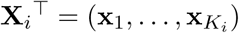 by

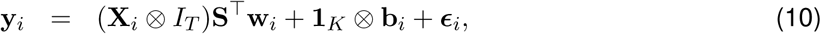

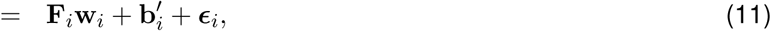

where 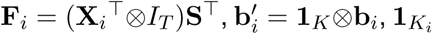 is a vector of 1’s of length *K*_*i*_, and ***ϵ***_*i*_^┬^ = (***ϵ***_*i*,1_^┬^, …, ***ϵ***_*i*_,*K*_*i*_ ^┬^). Equation (11) has the form of a standard multivariate linear regression. Therefore, if we set the noise distribution to be 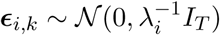 then we have the conditional distribution of **y**_*i*_ as

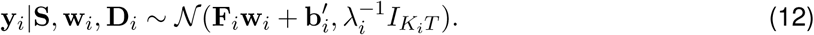

#### S1.1 Reduced inference in terms of S

Our strategy for accurate estimation is to focus on estimation of only one set of factors (**w**’s or **S**’s). In principle, either set of factors may be selected. In practice however the set of factors with lowest dimension should be selected to keep computational costs no higher than necessary. In the present case we have *T* ≪ *n* and we therefore will estimate **S**’s after integrating over **w**’s. In general, if we define a prior over **w**_*i*_’s denoted by *p*(**w**_*i*_) then the marginal likelihood of **s** is given by

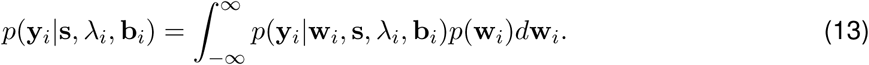

**Figure S1:**
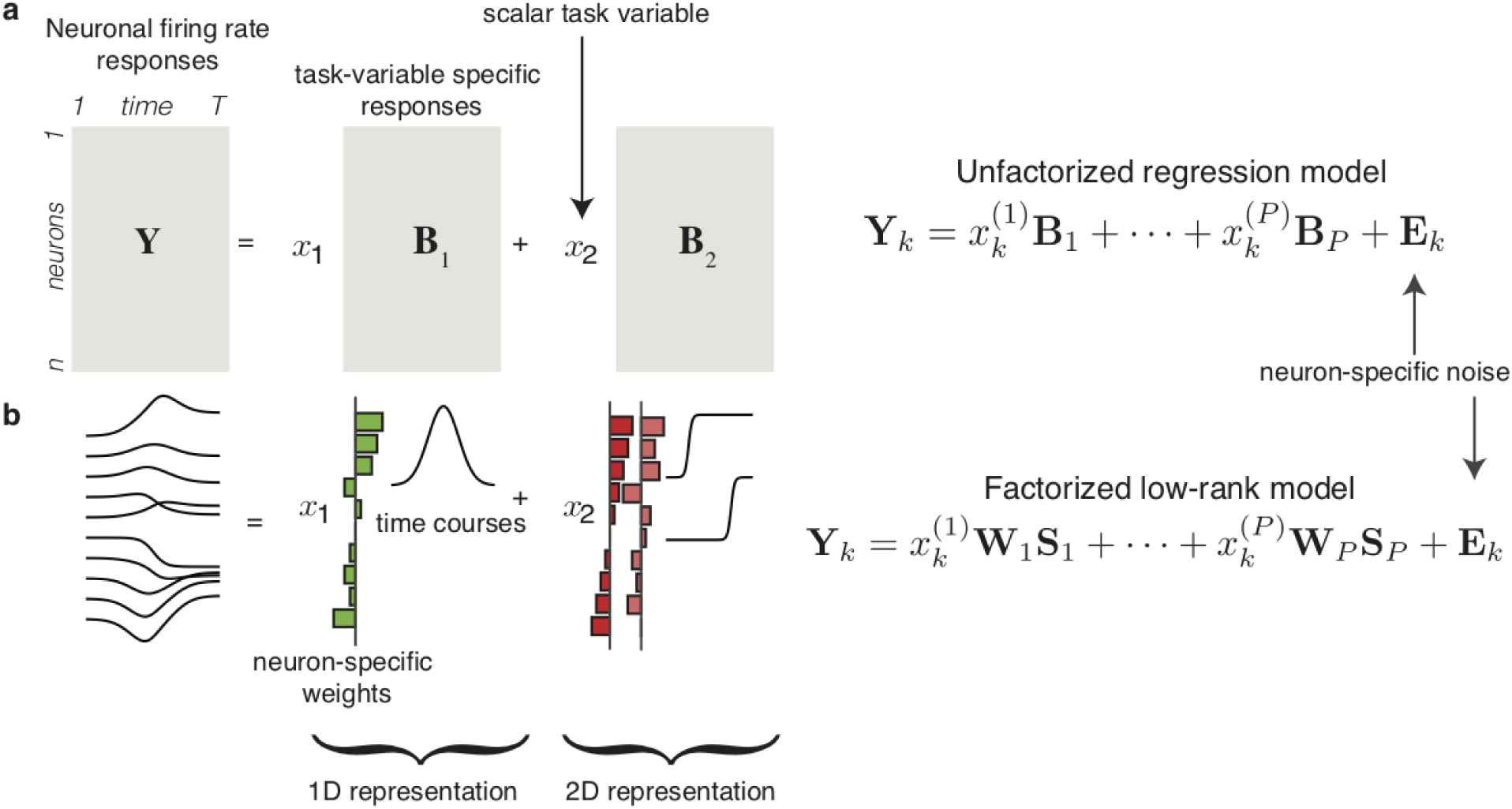
Schematic of low-rank structure in proposed regression model. **a)** The firing rates for *n* neurons observed over *T* time point on a give trial can be concatenated into a *n* × *T* matrix **Y**. A characteristic response for task variable and each neuron can also be described by a *n* ×*T* matrix **B**_*p*_ where the model linearly scales the characteristic response by the task variable. This formulation is equivalent to parameterizing each time point for each neuron as a separate linear regression problem. **b)** A low-dimensional description of the neural responses is achieved by parameterized each of the characteristic response matrices **B** by a small number of temporal basis functions. In the case of a 1D representation, a single temporal basis is needed, which is weighted separately to provide the response for each neuron. In the case of a 2D representation, two linearly-independent basis functions are needed, where each basis function gets its own set of weights to construct the characteristic responses of each individual neuron. The low-dimensional description is equivalent to a low-rank matrix factorization model for each **B**_*p*_.

If *p*(**w**_*i*_) is Gaussian then we can use standard Gaussian identities ^52^ to marginalize over **w**_*i*_ and obtain an analytical expression for the marginal likelihood in terms of **s**. In the present study, we let 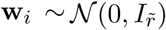, for all *i*, where 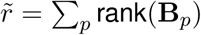. While this framework supports the application of a number of structured priors for *p*(**w**_*i*_), in the present work we utilize the conservative assumption of independent weights. While the scale of **w**_*i*_’s is inherently set by the prior, the scale of the 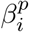 vectors will be learned by unconstrained estimation of **S**.

For the given likelihood and prior, our marginal likelihood is given by

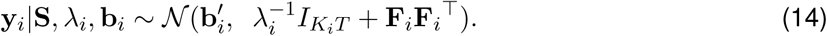

Assuming that noise correlations are negligible (in our case neurons are treated as having been recorded sequentially so that this is a reasonable assumption) we observe that neurons are conditionally independent given **S**. Thus, both the conditional and marginal distributions for the whole population factorize across neurons. Therefore, the population log-likelihood is given by

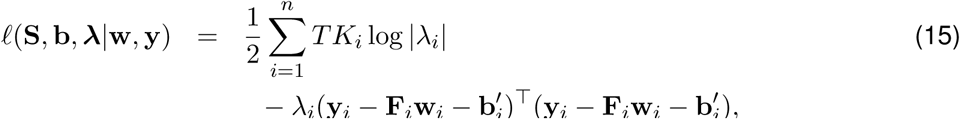

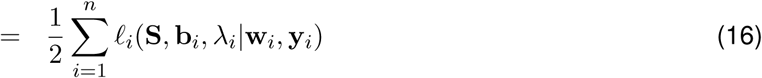

where *K*_*i*_ is the total number of trials observed for neuron *i*. The corresponding marginal log-likelihood is given by

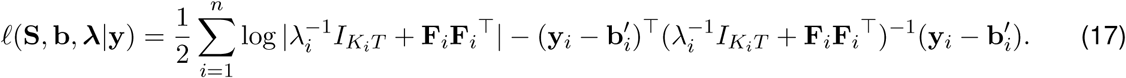

#### S1.2 Reduced expression for likelihood

It should be noted that the above marginal likelihood requires the log determinant and inverse of all *n* of the matrices 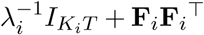 which are *K*_*i*_*T* × *K*_*i*_*T*. Thus, if all neurons are observed for *K* trials, then the determinant and inverse in general will have computational complexity *𝒪* (*nK*^3^*T* ^3^), which can be prohibitively large for even moderately sized datasets. Luckily, the expression for *ℓ* (**S, b**, **λ| y**) can be dramatically reduced.

After some algebra, we can derive the following expression for the marginal likelihood in terms of **S, λ**^┬^ = (λ_1_, …, λ_*n*_), and **b**:

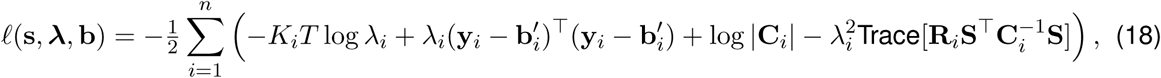

where

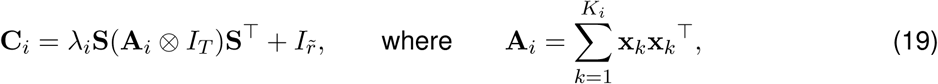

and the matrices **R**_*i*_ are defined by the outer product

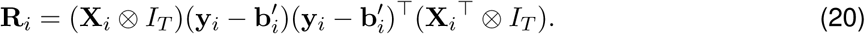

This formulation reduces the computational complexity to 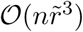 where, in general, 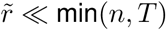.

#### S1.3 Posterior distribution w_*i*_’s

Common Gaussian identities can also be used to derive the posterior distribution over weights **w**_*i*_. As above, conditioned on **S**, the posterior distribution of all **w**_*i*_ factorize over neurons and we can write out each distribution independently. For the above model we find that

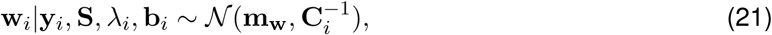

where

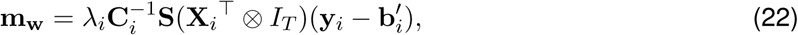

and **C**_*i*_ is defined in equation (19).

### S2 Parameter estimation

We estimate parameters by obtaining maximum likelihood estimates of **s, b**_*i*_, and **λ** by maximization of the marginal likelihood (18). The above description of the marginal likelihood *p*(**y**|**s, b, λ**), the complete data likelihood *p*(**y, w**|**s, b, λ**) = *p*(**y**|**w, s, b, λ**)*p*(**w**), and the posterior distribution over **w**, *p*(**w**|**y, b, s, λ**), allows us to derive an efficient algorithm for iteratively estimating **s, b**_*i*_, and **λ** using exclusively closed-form updates. The algorithm is essentially a special case of the “expectation-conditional maximization, either” algorithm (ECME) ^53^ where parameters are block-wise estimated by either maximizing the conditional expectation of the complete data log likelihood or the marginal likelihood.

#### S2.1 ECME for maximization of the marginal likelihood

Each iteration of our ECME algorithm comes with a conditional EM step (ECM) where we block-wise estimate **λ** and **s** while holding all other parameters fixed, followed by a direct maximization of the marginal likelihood in terms of **b**_*i*_. The E-step is defined by forming of the so-called “Q-function” for each neuron, which is an expectation over the complete data log likelihood and is given by

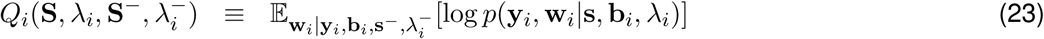

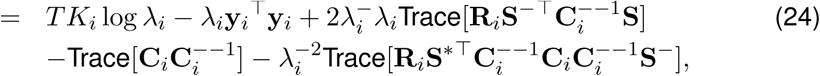

where 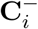 is as given in equation (19) except

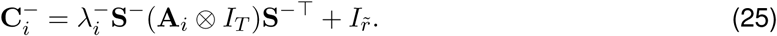

At each M-step we maximize *Q* ≡ ∑_*i*_ *Q*_*i*_ in terms of **λ** and **s** sequentially. Using updated estimates of **λ** and **s**, we then update **b** by direct maximization of the log likelihood (18). An outline of the algorithm is given in Algorithm 1.

The update for λ_*i*_ is obtained by setting *∂Q*_*i*_(**S**, λ_*i*_, **S**^−^, λ^−^)*/∂*λ_*i*_ = 0 and solving for λ_*i*_. The resulting update is given by

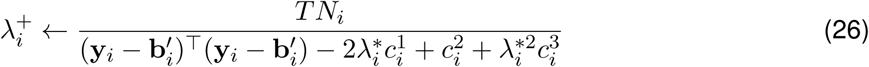

where

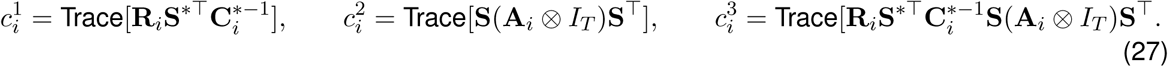

Similarly, the update for **S** is obtained by setting *∂Q*(**S, λ, S**^−^, **λ**^−^)*/∂***S** = 0 and solving for **S**. The

##### Algorithm 1 ECME for parameter estimation

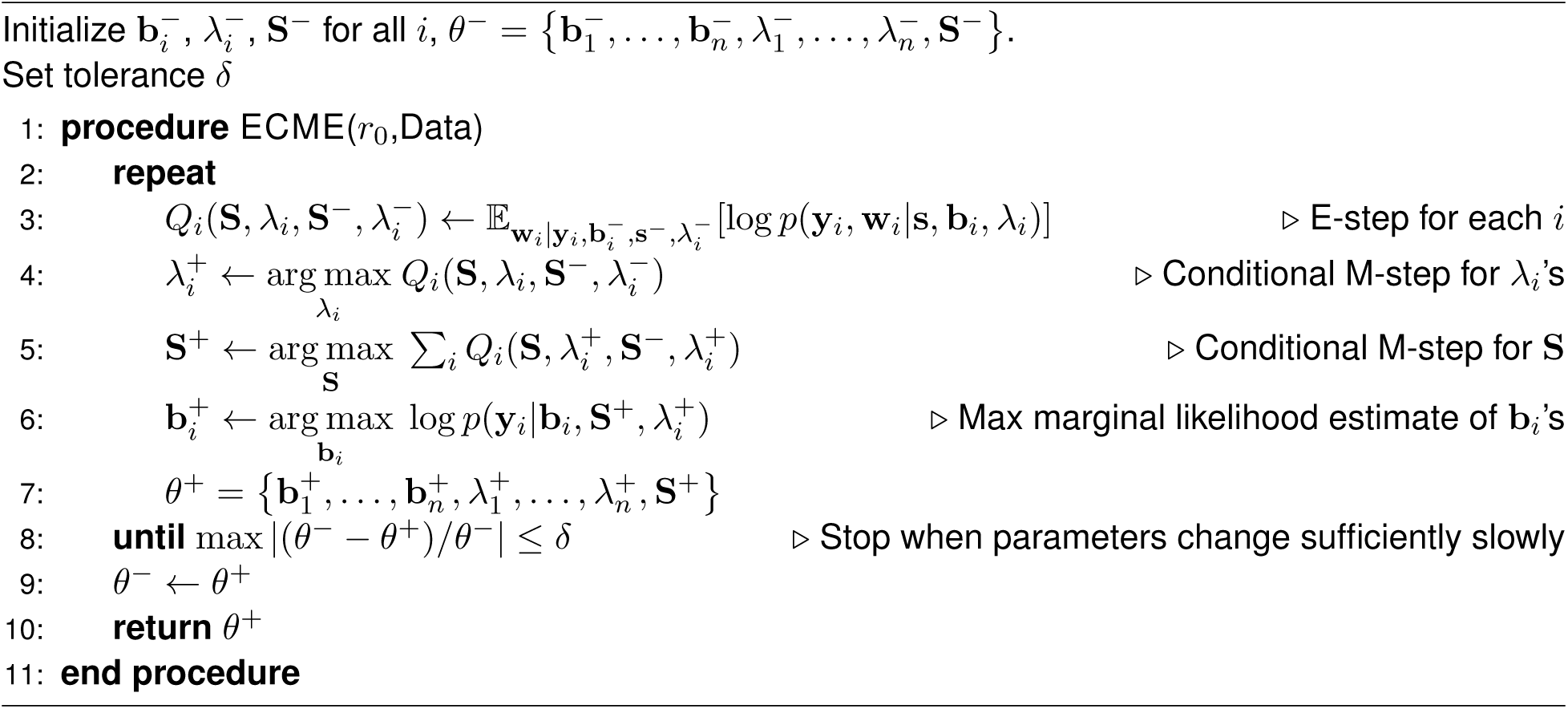

estimator satisfies the equation

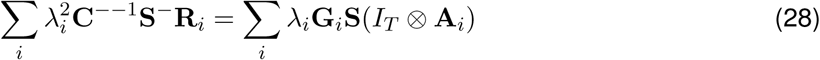

where

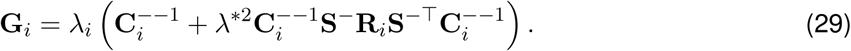

The solution to (28) is the solution to a linear system of equations that can be efficiently solved in 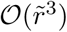 time.

Finally, the update for **b**_*i*_ is obtained by setting 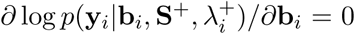 and solving for **b**_*i*_. The solution is given by

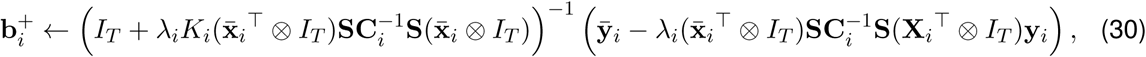

where 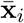 and 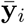 are the trial-averaged task variables and responses, respectively.

#### S2.2 Using ECME and direct maximization

While the EMCE method described above results in accurate estimates of parameters, the ECME method converges very slowly when it gets close to a local optimum. The problem of slow convergence is well documented among EM-type algorithms for related models like factor analysis ^53–55^. On the other hand, the ECME method gets close to a local optimum extremely fast.

Alternatively, we could directly maximize the marginal likelihood by gradient decent; an approach we will call maximum marginal likelihood estimation (MMLE). Although in principle the ECME method and the MMLE method should both be maximizing the marginal likelihood, they do so at different rates depending on distance from the optimum. In order to make best use of both methods we initialize using the ECME algorithm, which we parameterize with a liberal stopping criterion, and then complete the estimation procedure with MMLE. We observed this approach to provide faster convergence that either the MMLE or EMCE methods alone.

### S3 A greedy algorithm for rank estimation

While our model can identify low-dimensional subspaces of any dimension up to *D*_max_ = min {*n, T*}, the dimensionality of each subspace must be specified *a priori*. While we can use standard model selection techniques to compare the goodness of fit between models with alternative configurations, an exhaustive search would require searching over 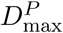 possible configurations. For our application this would mean estimating parameters for 15^7^ = 170, 859, 375 different models. We therefore developed a greedy algorithm for estimating the optimal dimensionality. A schematic illustration of the rank estimation procedure is depicted in Figure S2.

Recall that the dimensionality of each task-variable encoding corresponds to the rank of the factorization of the matrix of characteristic responses **B**_*p*_. We begin the algorithm by first estimating the model parameters with rank *r*_*p*_ = 1 for all *p*, giving us a model with total dimensionality 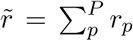 at the first iteration (i.e. 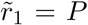) (Fig. S2, Iteration 0). At the *j*^*th*^ iteration we estimate the parameters of *P* models, each with the dimension of one of the task variables increased by 1, while keeping all other dimensionalities the same as in the previous iteration. We then get *P* models with total dimensionality 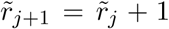 (Figure S2, Iteration 1-4). We then evaluate the AIC of each of these *P* models and keep the model with lowest AIC for the next iteration. In this way we grow the total dimensionality of the model by one on each iteration. The sample path that the algorithm produced for estimation of the ranks for data from monkey A is shown in Figure S3.

#### S3.1 Evaluation of dimension estimation with simulated data

Here we demonstrate that our rank estimation procedure recovers the true rank of the model the vast majority of the time even under conditions of vary small numbers of trials number relative to the size of observations. We also achieve good dimensionality estimation under model misspecification where the observations are drawn from a Poisson distribution (Fig. S4). We also examined the quality of rank estimates compared to alternative procedures for fitting the parameters and found that our method dramatically out-performed the alternatives.

We applied our greedy algorithm on simulated data in order to determine if it could accurately recover the true ranks using *n* = 100 neurons and *T* = 15 time points. For each run of our simulations we first selected a random dimensionality between 1-6 for each of *P* = 3 task variables (two graded variables with values drawn from {− 2, − 1, 0, 1, 2} and one binary task variable with values {− 1, 1}). Using these dimensionalities, the elements of **W**_*p*_ and **S**_*p*_ were drawn independently from a *𝒩* (0, 1) distribution. To give us heterogeneous noise variances, the noise variance for each neuron was drawn from an exponential distribution with mean parameter *σ*^2^ = 50. The resulting average SNR for any one task variable was −0.26 (±0.75, log10 units). We then simulated observations according to our model with varying numbers of trials (*N* ∈ {50, 200, 500, 1000, 1500, 2000}). In order to simulate incomplete observations, we set the probability of observing any given neuron on any given trial to *π*_obs_ = .4.

For each set of observations, we estimated the parameters of the model in one of three ways, which we describe below. In order to implement the AIC a likelihood and a degrees of freedom *K* must be specified. For all methods, on each iteration of the dimension estimation algorithm, we assumed a fixed dimensionality of the *p*-th characteristic response **B**_*p*_ to *r*_*p*_.

**Figure S2:**
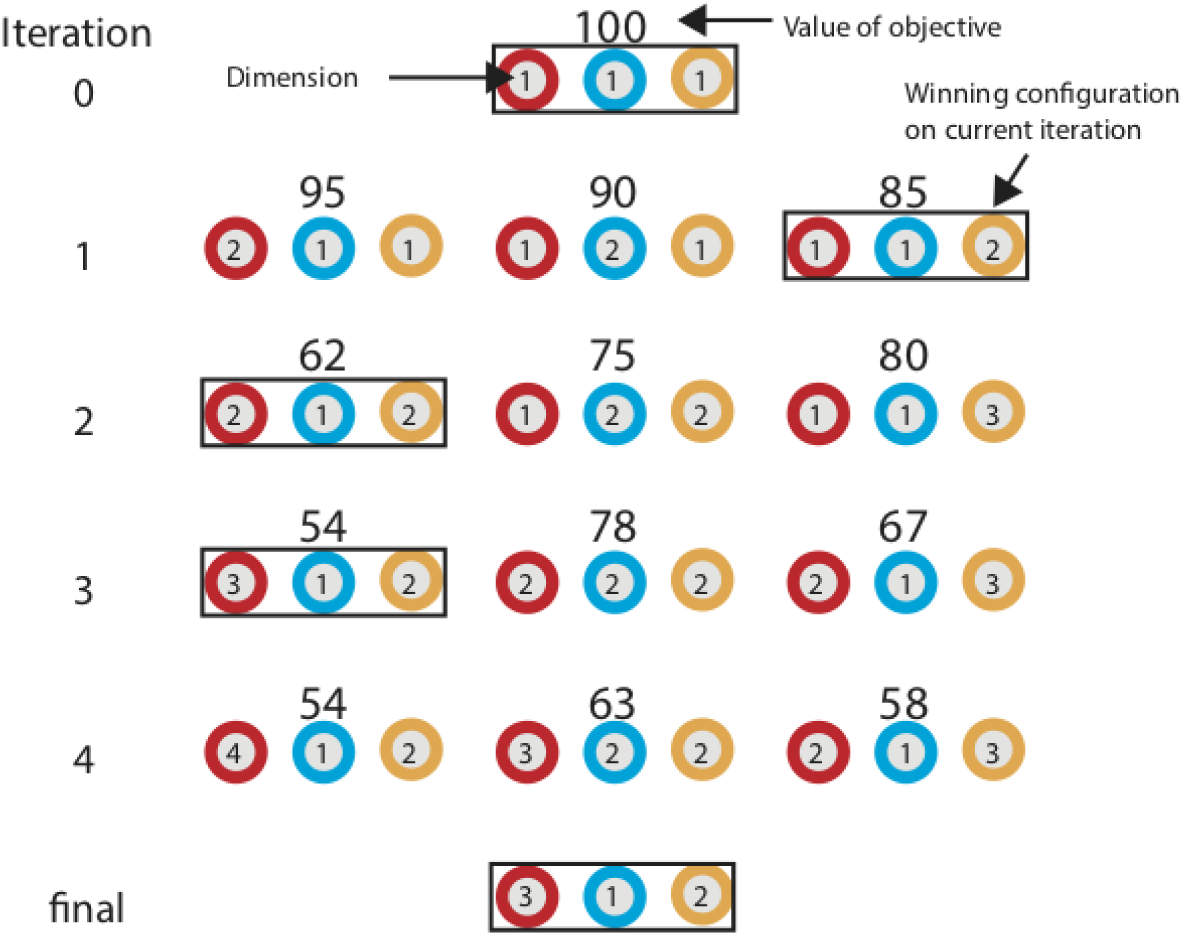
Graphical illustration of one possible sample path of algorithm for greedy estimation of dimensionality. Each colored circle represents a different task variable. Numbers inside of circles indicate dimensionality of the corresponding task variable at the current iteration. At iteration 0, the dimension of all task variables is set to 1. At the next iteration, all possible models with total dimensionality 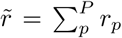 increased by 1 are evaluated by an objective function (in this case, the AIC). The configuration with the smallest value of the AIC will be selected as the starting point for the next iteration in which all possible models with 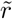 increased by 1 are evaluated. This process continues until the AIC cannot be decreased any further.

We considered the following four methods of parameter estimation:

1. *Linear regression and SVD* The elements of **B**_*p*_ for all *p* were estimated by linear regression for each neuron and time point independently. Each estimate of the complete matrix **B**_*p*_ could then be expressed by its singular value decomposition (SVD) as **B**_*p*_ = **U**_*p*_**D**_*p*_**V**_*p*_^┬^, where **D**_*p*_ is the *n* × *T* diagonal matrix of *d* = min{*n, T*} singular values. We then set the smallest *d* − *r*_*p*_ singular values to zero with the resulting matrix of *r*_*p*_ nonzero singular values denoted by 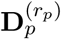. The rank-*r*_*p*_ estimates of **W**_p_ and **S** are then given by 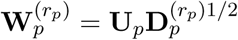 and 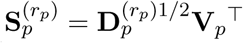, with the corresponding rank-*r*_*p*_ estimate of **B**_*p*_ given by 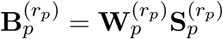. The corresponding likelihood is given by

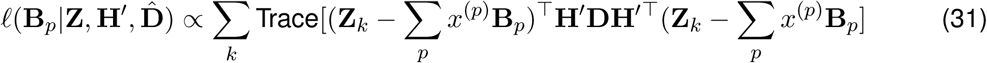
2. *Bilinear regression* After initializing with the rank-*r*_*p*_ estimates of **W**_*p*_ and **S**_*p*_ from the SVD method, the parameters can be further refined by bilinear regression. On each iteration, the values of **W**_*p*_’s are fixed, which leads to closed-form updates for conditional maximum likelihood estimates of **S**_*p*_’s and vice versa. Thus, the algorithm will alternate between estimating **W**_*p*_s and **S**_*p*_s until convergence. The bilinear regression method uses the same likelihood as (31).
3. *ECME* After initializing with the SVD solution we applied our ECME algorithm described in Section S2.
4. *Maximum marginal likelihood (MMLE)* After initializing with the ECME estimates of **W**_*p*_ and **S**_*p*_, we estimate **S**_*p*_ by direct maximization of the marginal likelihood given by (18). No estimation of the **W**_*p*_ factors is required since the marginal likelihood only depends on **S**_*p*_.

**Figure S3:**
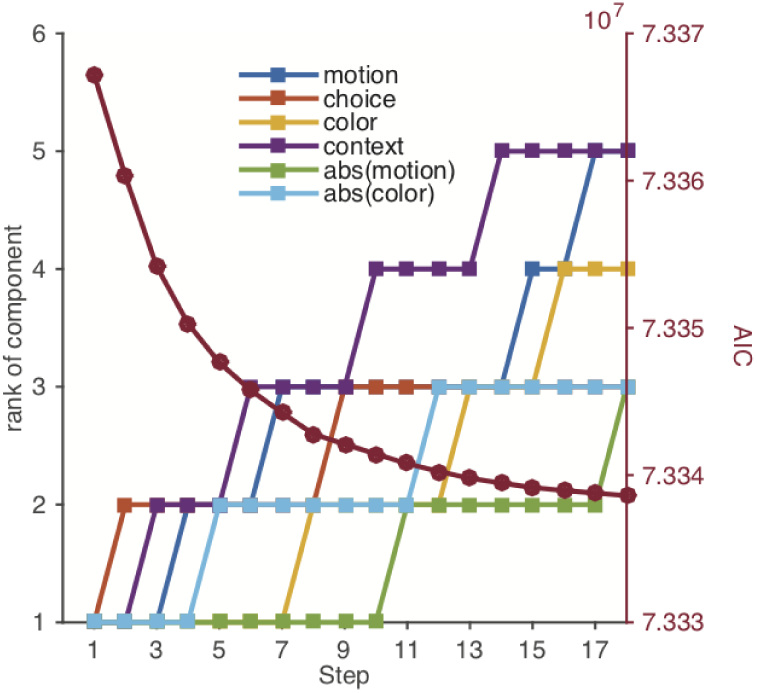
Sample path of rank estimation. On each iteration of the algorithm the total dimensionality of the model is increased by 1. Each color indicates the dimensionality of a different task variable after every iteration. The AIC on each iteration is shown in maroon.

For each setting of trial number *K*, we repeated this process 100 times and evaluated how well our algorithm estimated the dimension of the task variables by evaluating the difference between the true and estimated dimension of each task variable and counting the number of times that difference was observed. The results are presented in Figure S4. We found that all three methods tended to underestimate the dimensionality of the **B**_*p*_’s as the number of trials decreased but that this underestimation was least pronounced with our MML method, for which the vast majority of estimates resulted in the correct ranks even in the case of *K* = 50. Note that not only is this half the number of trials as neurons but since each neuron was only observed on about 40% of the trials this gives an average of 20 trials per neuron.

In order to evaluate the effects of model mismatch where the observations were drawn from a Poisson distribution, we repeated the above experiment using the MMLE method at *K* = 2000. We found no difference in the accuracy of the method between the case of Gaussian observations and the case where observations were Poisson (Fig. S4, dashed black line).

**Figure S4:**
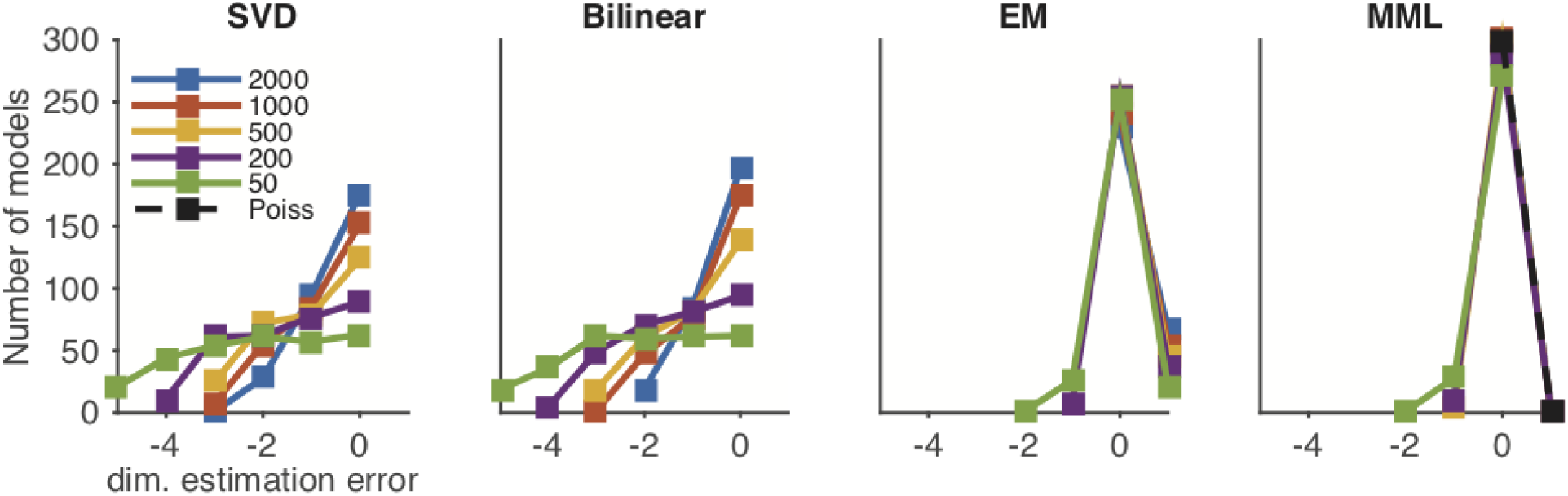
Simulation results for accuracy of rank estimation. Number of times (out of 100) that the difference between the estimated and true dimensionality (dim_est_ - dim_true_) of each of the *P* = 3 characteristic response matrices **B**_*p*_, giving a max count of 300.

### S4 Specifying subspaces

#### S4.1 Subspace Identifiability

We note that the factorization **B**_*p*_ = **W**_*p*_**S**_*p*_ is not unique and leaves the model parameters only identifiable up to rotation and scalar multiplication. Specifically, note that we can define a orthonormal rotation matrix **P** and a scalar *α* to obtain a new pair of matrices 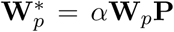 and 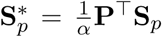 such that 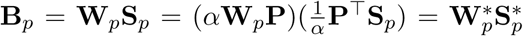. This non-identifiability is identical to the type of non-identifiability inherent to other matrix factorization models such as factor analysis or probabilistic PCA ^56^. Therefore, we require a way of uniquely identifying the subspace spanned by **W**_*p*_.

We can obtain a fully identifiable subspace by first reconstructing **B**_*p*_ from the estimated **S**_*p*_, where the **W**_*p*_ is estimated from the expectation of the posterior of **W**_*p*_ given in (22). Each **B**_*p*_ will then have a unique singular-value decomposition (SVD) denoted by **B**_*p*_ = *U*_*p*_∑_*p*_*V*_*p*_^┬^. We then take the first *r*_*p*_ columns of *U*_*p*_, denoted 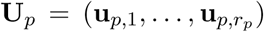, to define the encoding subspace of task variable *p* where we will refer to the *j*th vector in this subspace as **u**_*p,j*_. In this way, we obtain an orthonormal basis whose orientation gives an ordered set of vectors where the order is with respect to the variance of **B**_*p*_ explained. We refer to this orientation as the *principle components* (PC) orientation due to its relation to principle components analysis.

#### S4.2 Orthogonalization of Subspaces

The mTDR model does not impose any orthogonality between task variables or task variable sub-spaces. This permits accurate recovery of subspaces even when the encoding dimensions are correlated, as we demonstrate in Supplementary section S8. It is desirable therefore to be able to visualize the part of the encoding of each task variable that is unmixed ^27^. We therefore orthogonalize the sub-spaces with respect to correlated subspaces.

To do this we first obtain the PC axes **U**_*p*_ defined in Supplementary section S4.1. Orthogonalization of the basis **U**_*p*_ with respect to some other set of basis vectors **U**_*q*_ was achieved by the Graham-Schmidt orthogonalization. For example, if we wished to orthogonalize a stimulus subspace with respect to the choice subspace, we form the concatenated matrix [**U**_choice_**U**_stim_] and orthogonalize to obtain the orthogonalized basis [**P**_choice_**P**_stim_] as in

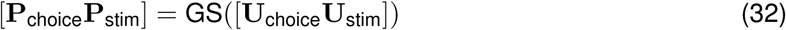

where GS(*M*) indicates performing Graham-Schmidt on the matrix *M*. Thus, **P**_stim_ (where “stim” = color or motion), is a set of orthonormal vectors that define the part of the stimulus subspace defined by **U**_stim_ that is orthogonal to **U**_choice_.

### S5 Projections onto jPCA axes

The low dimensional projections in Figure 4 exhibit rotation-like dynamics. In order to verify the rotational nature of these projections and identify the plane of most rotation-like dynamics, we used jPCA ^18^ (calculated using Matlab code obtained from http://stat.columbia.edu/cunningham/). Projections onto the first two jPCA axes are presented in Figure S10.

In order to examine whether or not rotational structure was trivially present in our data we first examined projections of shuffled versions of the data. Each neuron’s PSTHs were shuffled with respect to trial type and projected onto the learned task variables axes. No clear sequential or rotational structure is observable (Figure S16). We performed jPCA on these projections and similarly found no qualitative evidence for rotations (Fig. S17).

To test for the presence of rotations more rigorously, we used a sampling method developed by Elsayed and Cunningham ^29^ in which we drew 100 samples from the maximum entropy distribution with the same second order moments as the data. We then learned a low-rank model for each sample, identified low-dimensional projections, learned a basis for the jPCA plane, and projected held-out trials onto this plane. From these projections we identified the angle of rotation and constructed a confidence interval (shown by the shaded regions in Fig. 4f).

### S6 Decoding

#### S6.1 Unconditional decoding

Once estimates of **B**_*p*_ and ***λ*** and obtained we can decode new trials using maximum likelihood. Because most neurons were not observed simultaneously, the specification of our observations **Y**_*k*_ in terms of the full set of neurons is incomplete. We accommodate non-sequential observations by specifying the true observations on each trial by **Z**_*k*_ = **H**_*k*_**Y**_*k*_ where **H**_*k*_ is an observation matrix. Suppose *n*_*k*_ *< n* neurons were observed on trial *k*, then **H**_*k*_ is a *n*_*k*_ × *n* matrix where each row is a “one hot” vector indicating that the corresponding neuron was observed.

If **Z**\*, **H**\* and **x**\* are new observations of the population response, observation matrix, and task variables, then the likelihood of **x**\*, conditional on 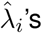, and 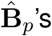 is given by the data log likelihood defined by (14), which will be proportional to

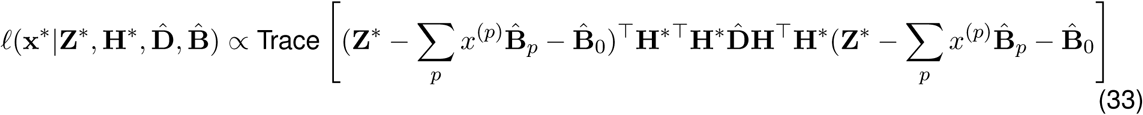

where 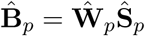.

Differentiating with respect to *x*^(*p*)^ gives

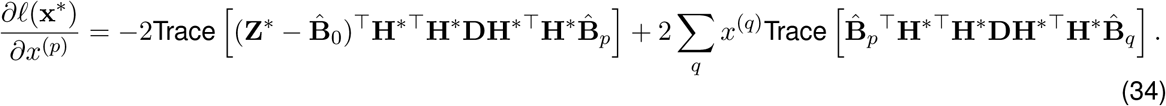

If we let **M**\* (*I*_*T*_⊗**D**^1*/*2^**H**\*^⊤^**H**\*) (vec(**B**_1_), …, vec(**B**_*P*_) and 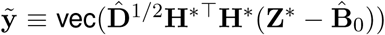, then we can write the gradient of *ℓ* (x) in vector form as

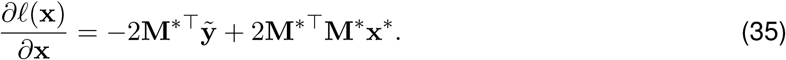

Setting 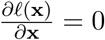 therefore, yields a closed-form solution for the maximum likelihood estimator for **x**\*,

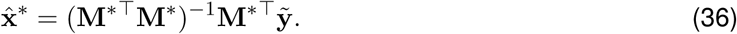

This formula is intuitive as we can see that 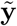 is a precision-weighted vector of the new observations, 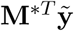 is the projection of these observations onto each of the estimated task variable subspaces, and (**M**\*^┬^ **M**\*)^−1^ serves to whiten the projection, accounting for the fact that the estimated subspaces are not necessarily orthogonal. The decoding weights are defined as (**M**\*^⊤^**M**\*)^−1^**M**\*^⊤^**D**^1*/*2^.

Instantaneous estimates of **x**\* at time *t* can be obtained by simply restricting **B**_*p*_ and **Z**\* to their *t*^th^ columns and following the same inference procedure.

#### S6.2 Conditional decoding

If we want to consider some elements of **x**\* to be known, then there is a straight forward way to do so. This may be the case, for example, when maximizing the log likelihood, conditioned on the animal’s choice when evaluating the log likelihood ratios.

Suppose we let task variables *p* = 1, …, *q* be unknown and task variables *p* = *q* + 1, …, *P* be known, and let x_1_ *≡* (*x*_1_, …, *x*_*q*_)^⊤^ and x_2_ *≡* (*x*_*q*+1_, …, *x*_*P*_) ^⊤^. Furthermore, we can define matrices 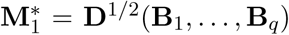 and 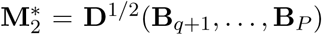. The maximum likelihood estimator for x_1_, conditioned on x_2_ is then given by

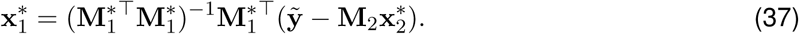

#### S6.3 Decoding of discrete variables by log likelihood ratio

The task variables in these data are a combination of discrete (choice, context) and continuous (color, motion) variables. It is therefore prudent to respect the domain of the discrete variable when decoding (*x*_*p*_ ∈ {1, − 1}). For example, when we decode for choice, we first calculate the MLE of the continuous variables, conditioned on the two possible choices (see Supplementary Section S6.2). This results in two vectors of task variable estimates (x^+^, x^−^), one for each choice. We then evaluate the log-likelihood at each of these vectors to calculate the log-likelihood ratio (LLR), which measures the relative information in favor of the two possible categories. For data given by **Z**\*, the LLR is given by

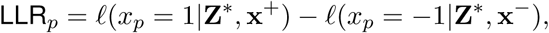

where *ℓ* (*x*_*p*_ = 1|**Z**\*, x^+^) is the log likelihood evaluated at *x*_*p*_ = 1, and x^+^ is the MLE of all other task variables, conditioned on *x*_*p*_ = 1. The inferred probability on a given trial that the data were generated with *x*_*p*_ = 1 is therefore given by

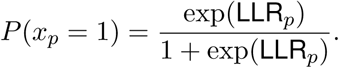

The value of this approach to decoding is that we obtain a probability of a trial category at each time point, and not just a candidate category, conditioned on the neural activity. Evaluating the likelihoods with the conditional MLEs (x^+^, x^−^) allows us to account for the confounding effects of the other task variables. The LLRs for context were calculated in an analogous way.

For Figures 7, S21, S22, S20 we evaluated the log likelihoods with the MLE of the stimulus estimates, conditioned on the corresponding discrete variable.

### S7 Interpretation of projection vectors

In order to draw principled connections between the projected PSTH’s and the decoded values of task variables, we adopted the following conventions for projections. We will carefully consider equation (37) and assume that the time-dependent (i.e. task-variable independent) component is given by **B**_*P*_. For simplicity, let us assume that all neurons have been observed.

First, recall from our description above on unconditional decoding that the projection of the data onto the subspace of unknown task variables at time *t* is given first by the projection of the normalized quantity

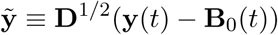

onto the regression weights as in

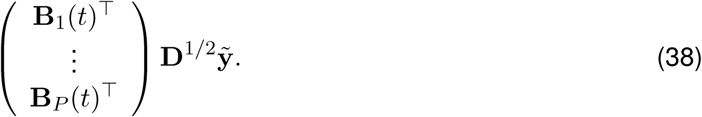

We can write this same expression along with the decomposition of **B**_*p*_, which is given by

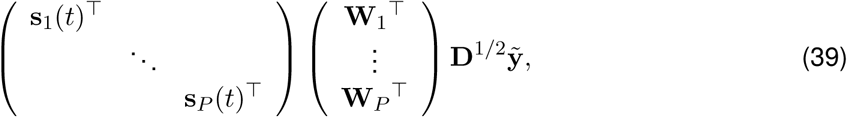

where s_*p*_(*t*) is a length-*r*_*p*_ vector corresponding to the collection of all *r*_*p*_ basis functions for task-variable encoding *P* at time *t*.

Therefore, there are two projections that take place to convert the mean-subtracted data into time-varying predictions of task variables. The first takes place by projecting the data onto the subspace defined by (**W**_1_, …, **W**_*P*_)^⊤^**D**^1*/*2^, which does not change with respect to time and preserves the dimensionality of encoding. The second projection is onto blkdiag(s_1_(*t*)^⊤^, …, s_*P*_ (*t*)^⊤^), which changes over time and reduces the dimensionality from 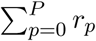 to *P*. Since the encoding subspace should be independent of time we therefore defined the low-dimensional trajectories by

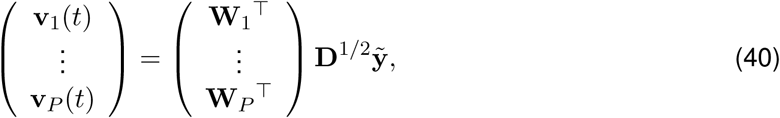

where 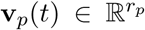 s the low-dimensional trajectory for task variable *p*. Rotations of these projections, such as those plotted using seqPCA were obtained by first identifying the rotation matrix **R**_*p*_ and projecting onto the rotation as in **R**_*p*_**v**_*p*_(*t*).

Therefore, decoding by maximum likelihood (Supplementary section S6) requires a linear transformation of the low dimensional trajectories **v**_*p*_(*t*). Specifically, the decoded task variables x*(*t*) are given by

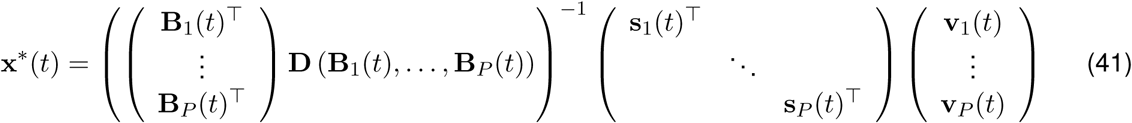

Therefore, because the decoding weights vary in time any variation associated with the encoding can be counteracted by the decoding.

### S8 Relationship between subspaces

We investigated the degree to which the characteristic responses reflected coordinated activity in two ways. First, we examined the subspaces correlations and second, we examined the degree of agreement between the subspaces thenselves using cannonical correlations analysis (CCA).

#### S8.1 Subspace correlations

Subspace correlations were calculated by taking the cross-correlations between characteristic responses (**B**_*p*_). Correlated responses imply that the population does not encode task variables independently and the encoding of task variables occurs in a (at least partially) shared subspace.

We examined the cross correlation between characteristic responses of task variables to visualize the change in correlations over time. The results of this analysis are displayed in Figure S5.

#### S8.2 Subspace agreement

We analyzed the overlap between task variable subspaces by performing CCA. We used CCA because it allowed us to identify alignment between subspaces that do not have the same dimensionality. The result is a sequence of correlation coefficients that describe mutually orthogonal directions where the subspaces are at least partially aligned. The results of this analysis are presented in Figure S6.

Signifiant, multi-dimensional overlap for both monkeys were observed between the motion-choice and color-choice subspace pairs. Smaller, but still significant overlap was also observed for motion-color, abs(color)-context, and abs(color)-abs(motion), subspace pairs. Monkey F showed stronger correlations across subspaces than monkey A. Monkey A showed no overlap between the abs(mo) subspace and the motion, color, or choice subspaces.

**Figure S5:**
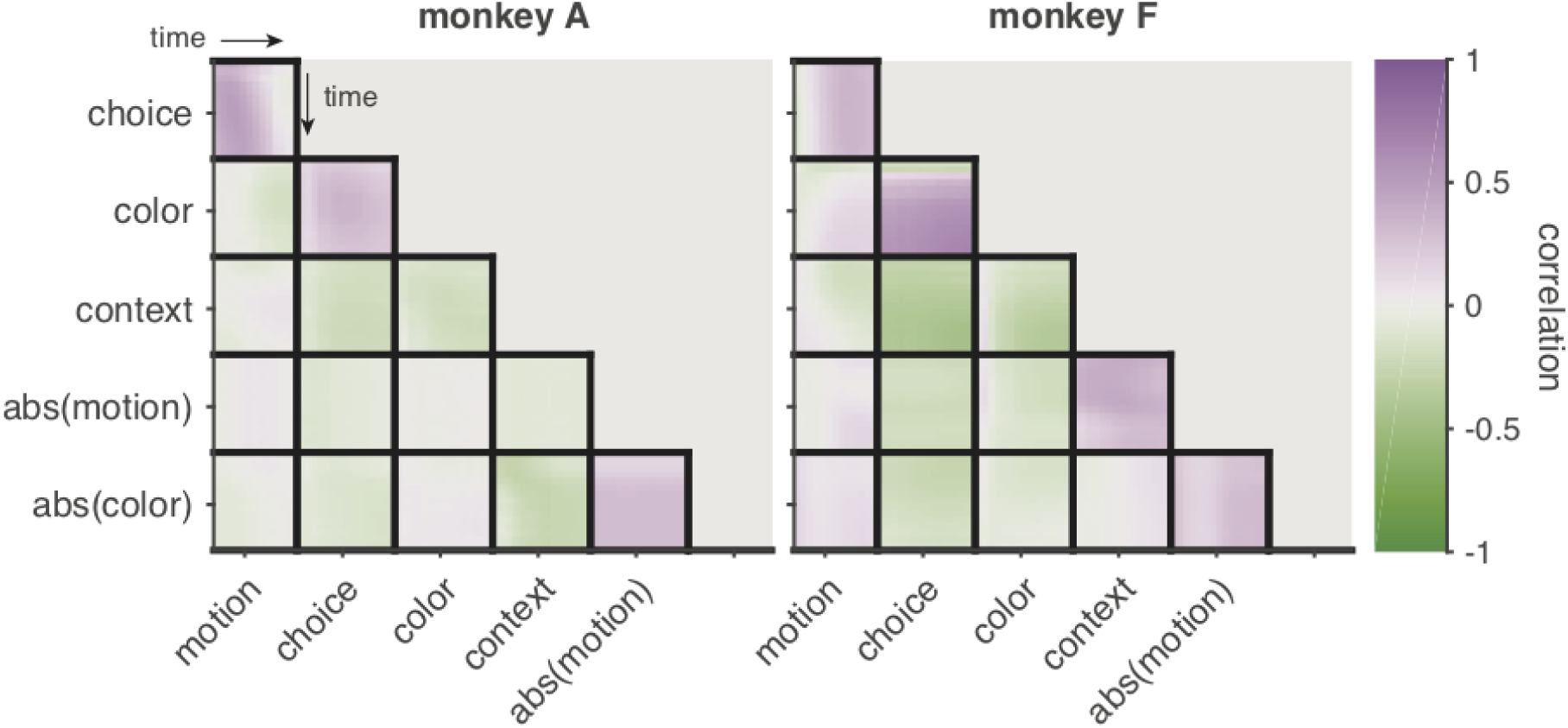
Cross correlation between characteristic responses of task variables. Motion and color coherence encoding appears to be positively correlated with the choice encoding for both animals.

### S9 Sequential PCA (seqPCA)

The goal of seqPCA is to identify a subspace orientation that best describes the data via a sequence of axes in which the order of the axes describes the order in time that each axis dominates the variance of the data.

The basis for seqPCA is constructed as follows: Suppose we have *D*-dimensional observations **y**_*t*_ at each time *t*. We can arrange all of the observations up to time *t* into a *D* × *t* matrix 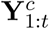 where the index *c* may refer to trials or conditions. We can arrange the data for all *c* = 1, …, *C*, up to time *t* into a *D* × *tC* matrix 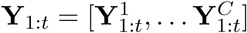. If the *j*^*th*^ singular value of **Y**_1:*t*_ is denoted by *σ*_*j,t*_ then the fraction of variance that the *j*^*th*^ singular vector describes for the first *t* time points is given by

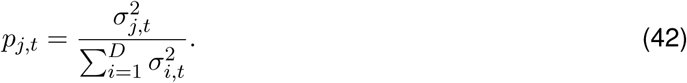

If the singular values are ordered such that *σ*_1,*t*_≥ *σ*_2,*t*_≥ … ≥*σ*_*D,t*_, then the largest possible variance captured by any single linear dimension at time *t* is given by *p*_1,*t*_.

This construction evokes a sequence of proportions such that *p*_1,*t*_ will vary in characteristic ways according to the specific dynamics of the data. For example, if the data project perfectly onto a single dimension then *p*_1,*t*_ = 1 for all *t*. If the data are unstructured then *p*_1,*t*_ will decrease monotonically until it converges to *p*_1,*t*_ = 1*/D*. However, if the data are structured such that the sequential observations progress linearly along a single direction up to time *t*′and then change direction, then *p*_1,*t*_ will increase up to time *t*′ and then begin to decrease.

**Figure S6:**
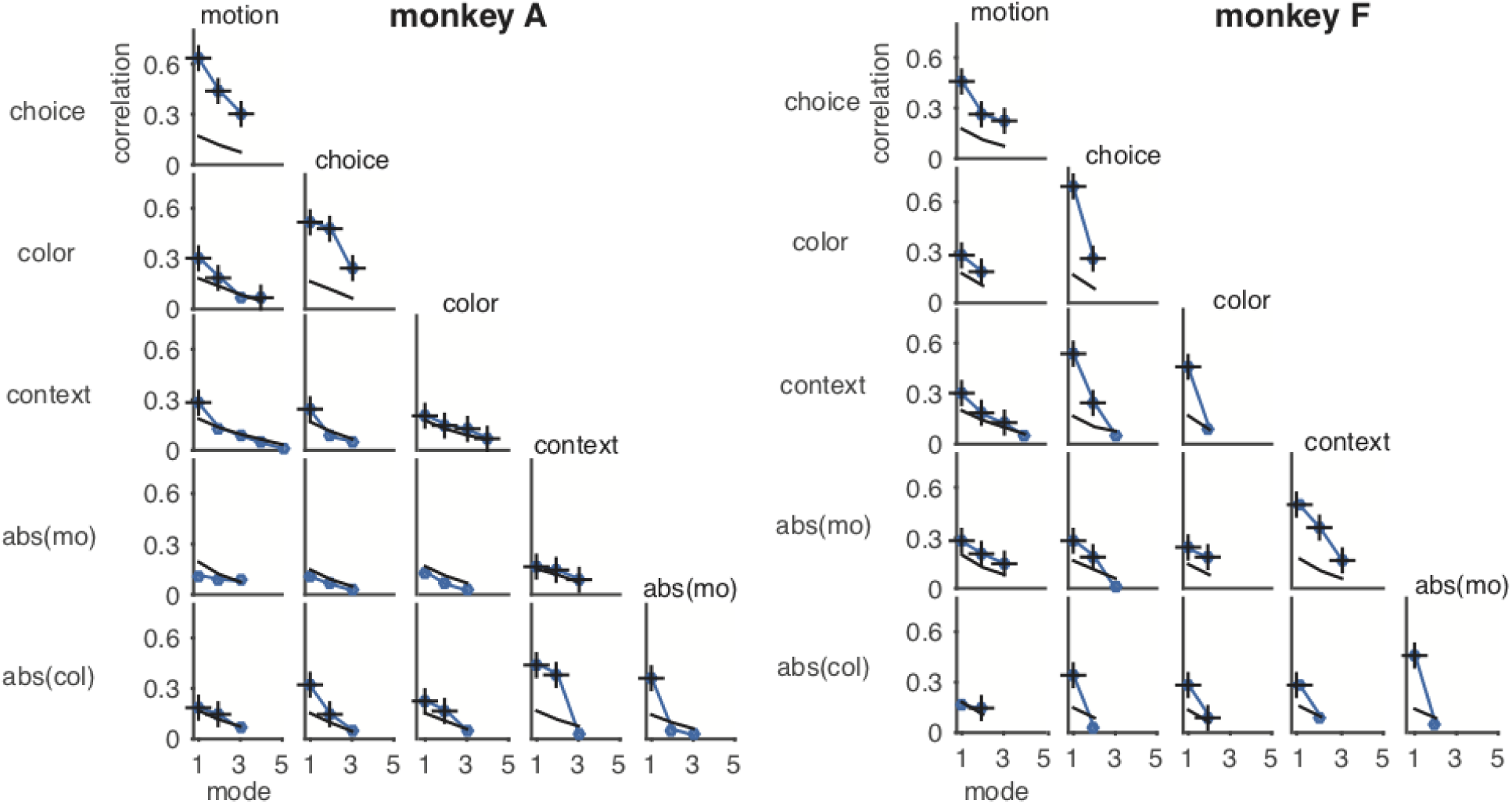
Canonical correlations between task variable subspaces. Canonical correlations measure the degree to which the subspaces overlap. Black lines indicate 95% confidence limit for canonical correlations from 100 randomly permuted axes from the measured subspaces. Markers with “+” indicate the measured canonical correlations that are significantly larger than expected by chance (permutation test, controlling for false discovery rate at .01 level).

In the latter case we can identify the point at which the data begin to change direction as a peak in the *p*_1,*t*_ sequence. If *t*′is the time of this peak then we can identify the first basis vector as the first singular vector at time *t*′ (**u**_1,*t*′_). To identify the next sequential element to this basis we can subtract off the projection onto the first basis as in

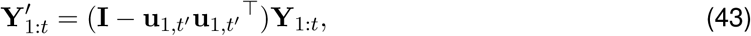

and repeat the process on 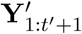. The process may be repeated *D* − 1 times with the *D*^*th*^ vector being completely determined. An orthonormal basis can be constructed from this collection of vectors by Graham-Schmidt orthogonalization.

While the method will always produce a basis for the data, the data never needs to load exclusively onto a single axis at each time point. This is a crucial detail in that exclusive loading onto a single seqPCA axis is a feature of the data, not the method.

We will illustrate the seqPCA method with the following 3D example. A sequential data set was generated by first generating 3 orthogonal vectors that were added. The coordinates of 10 points uniformly spaced on each line were jittered by adding Gaussian noise (Fig. S7a).

The seqPCA algorithm starts by calculating the variance explained by the first singular vector. As the number of data points increases, the first singular vector explains more of the variance until it reaches the 10th data point, after which it decreases, followed a a second, smaller peak (Fig S7b, left). This first peak represents the point at which the data matrix includes all of the first ten data points, shown as the blue points in Fig. S7a. After these first 10 points all other points necessarily lie in an orthogonal subspace and the amount of variance explained by any one axis necessarily decreases. Therefore, the first peak represents the number of datapoints to include to calculated the first seqPCA axies (seqPC_1_), represented by the blue line in Fig. S7a. The index of this peak serves as the temporal boundary between the seqPC_1_ and seqPC_2_ axes.

After the seqPC_1_ has been identified, the projection of the data onto seqPC_1_ is subtracted from the data as described in (43) and the algorithm picks up the analysis using the residuals from the first temporal boundary. We find a second peak around time index 20 (Fig. S7b, right). This peak correctly identifies the transition between orange and yellow data points in Fig. S7a.

**Figure S7:**
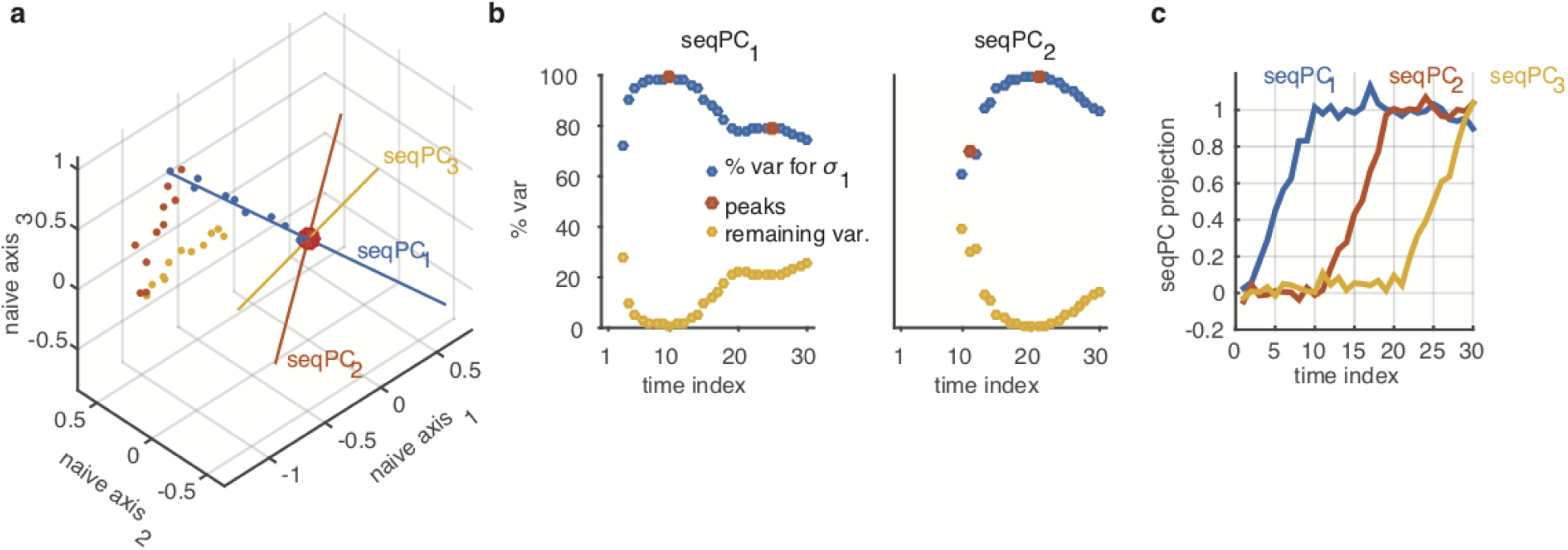
**a)** Example data (dots) and estimated seqPCA axes (colored axes). **b**) Example of seqPCA vector selection process using motion subspace projections. Blue markers indicate the fraction of variance explained by the first left singular vector (*p*_1,*t*_), compared to all remaining dimensions, at each time index. **c**) Projection of data onto the estimated seqPC’s.

### S10 Relationship between early/middle/late axes and TDR axes

We compared projections obtained through mTDR and the TDR method proposed previously ^1^. While the steps that lead to acquiring a projection axis in the two methods differ substantially, most of these steps are aimed at denoising and regressing the data. The key features of each method is in the selection of the subspace to be analyzed once regression weights have been identified. The TDR analysis chose a single axis for each subspace, corresponding to the regression coefficients at the time index with maximum norm, and then performed an ordered orthogonalization of these axes. Formally, if **B**_*p*_(*t*) is the vector of regression coefficients for task variable *p* at time index *t*, then the non-orthogonalized axes are identified by

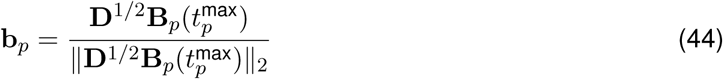

where

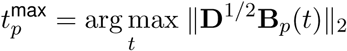

and **D** = diag(***λ***) is the diagonal matrix of noise precisions (see Supplementary Section S1). The axes are orthogonalized by first arranging the vectors into a matrix as [**b**_choice_ **b**_motion_ **b**_color_ **b**_context_] and then orthogonalized by the Graham-Schmidt algorithm. We normalize the regression coefficients by **D**^1*/*2^ to reflect the fact that the neurons were Z-scored prior to regression in the previous analysis ^1^.

We obtained projection weights for the mTDR method first by identifying the low-rank matrices of regression coefficients by maximum likelihood (ML) as described in previous sections of this supplement (Sections S2, S4.2, S7, and S9), performed an ad hoc orthogonalization on the stimulus and context subspaces (see caption of Fig.4) and then rotated them (ML + rotation) to obtain the early, middle, and late seqPCA axes for each subspace.

#### S10.1 1D TDR versus multidimensional mTDR and projection magnitudes

Encoding magnitudes were compared (Fig. 4e, S12) by comparing the projections obtained from the TDR encoding axes (Supplementary noteS10) with those of the mTDR method where the mTDR projection was summed across early, middle, and late axes. This is appropriate since the three seqPCA axes are orthogonal to each other. While Figures 4e, S12 only display the strongest encoding strengths, statistical testing was conducted using pseudotrials drawn for all stimulus strengths. Paired, left-tailed Wilcoxon signed-rank test was used to test whether mTDR more strongly encoded (i.e. projections are further from zero) that those of TDR. The positive false discovery rate ^28^ (pFDR, controlled at .01) was used to control for multiple comparisons.

#### S10.2 Correlations between TDR and mTDR axes

We examined the correlation (i.e. the normalized inner product) between the early/middle/late axes and the axis selected by the TDR max-norm approach described by equation (44). The correlations are presented graphically in Fig.S8. Figure S8 shows that while the TDR axes are weakly correlated with all three seqPCA axes, they are best aligned with the early axis, quantitatively confirming the qualitative similarity between the trajectories from previous TDR analysis and the trajectories presented in Fig. 4.

**Figure S8:**
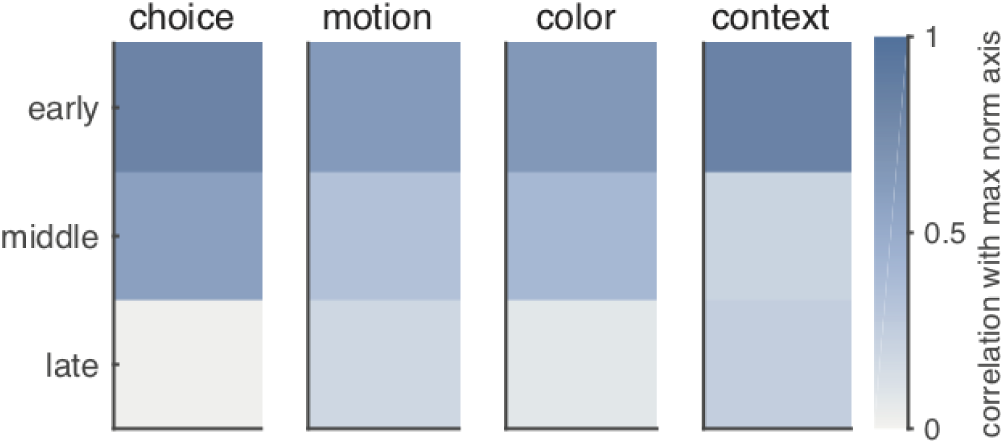
Correlations between max-norm axes and early/middle/late axes for each subspace. For all subspaces the maximum correlation between the max-norm axis and the early axis is larger than the correlation between the max-norm axis and the middle and late axes.

## S11 Supplementary figures

**Figure S9:**
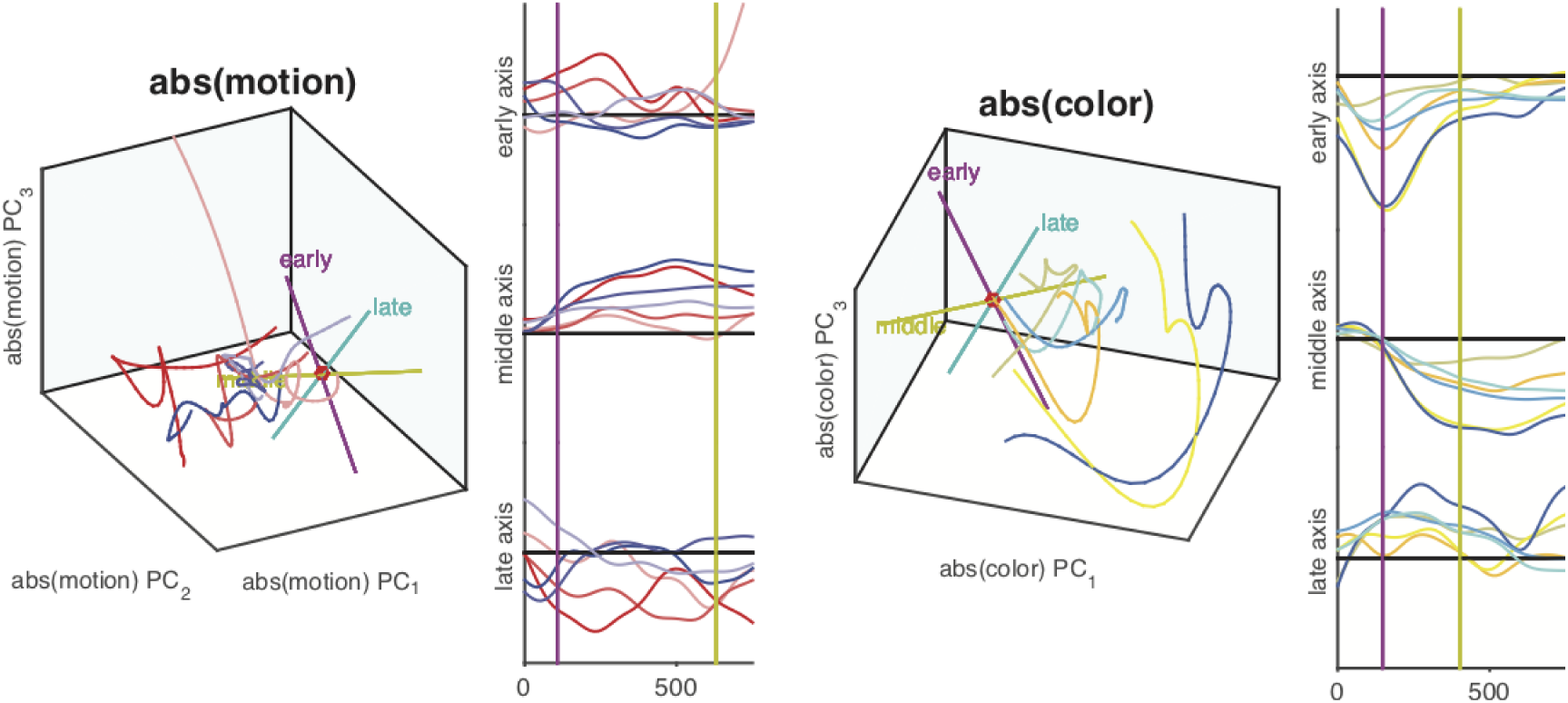
Projections of population PSTH’s onto the first, second, and third PC-axes for monkey A. **a)**The abs(motion) and **b)** abs(color) subspaces. Subspaces have been orthogonalized with respect to the first dimension of the choice subspace. The monkey gave the correct response for all trials used. Colored axes indicate dominant axes in the early, middle, and late periods of the stimulus epoch, as determined by the methods described in Supplementary section S9. Purple vertical lines indicate transition from the early to middle epochs. Yellow vertical lines indicate transition from the middle to late epochs as in Figure 4. Plotting colors are the same as those in Figure 4. Units of the ordinate are arbitrary but all axes are on the same scale.

**Figure S10:**
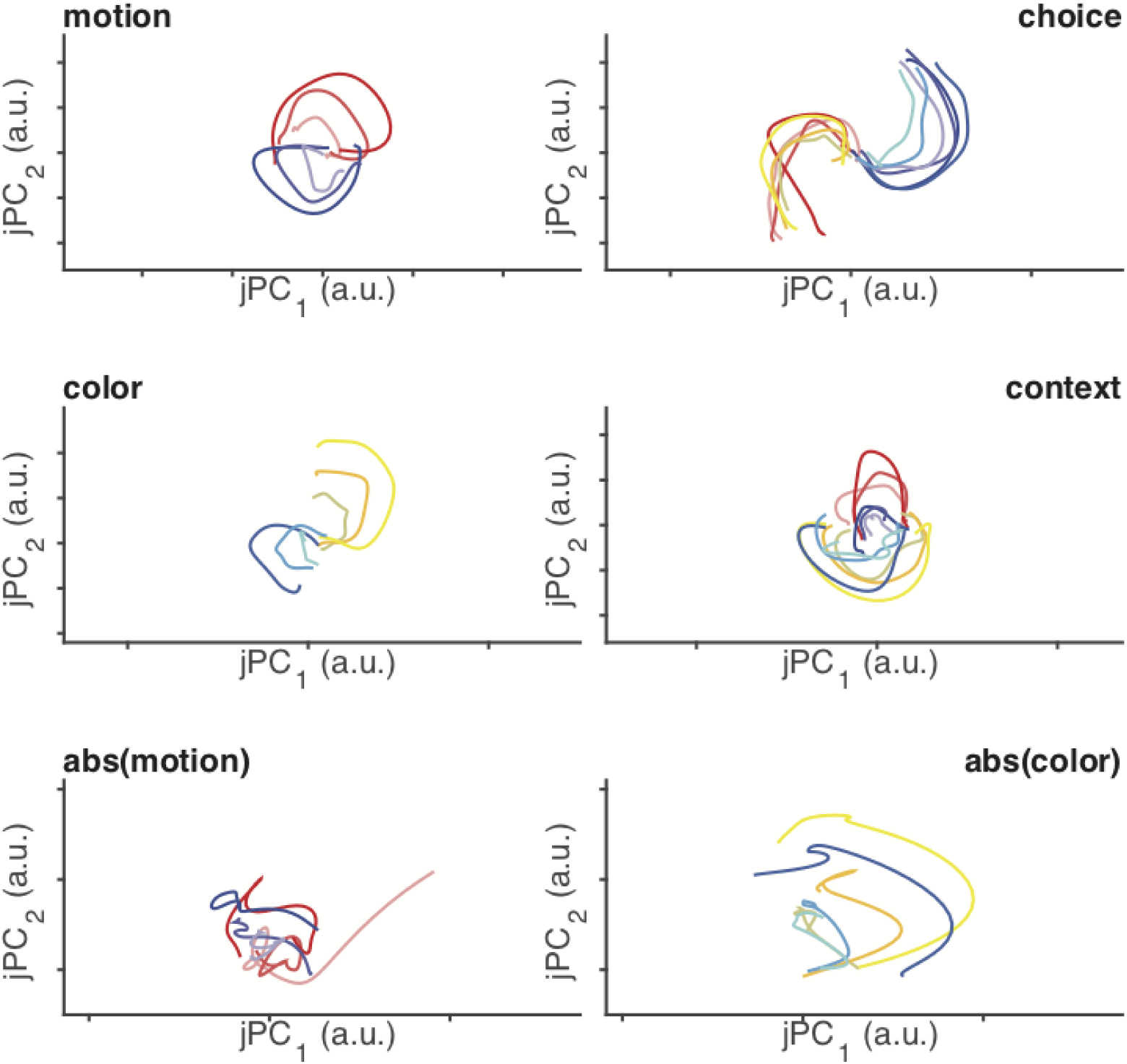
Projections of population PSTH’s onto jPCA axes for monkey A. Projections are onto the first two jPCA axes identified by the trajectories shown in Figure 4. The jPCA axes reveal strongly rotational dynamics for motion, color, choice, and context subspaces.

**Figure S11:**
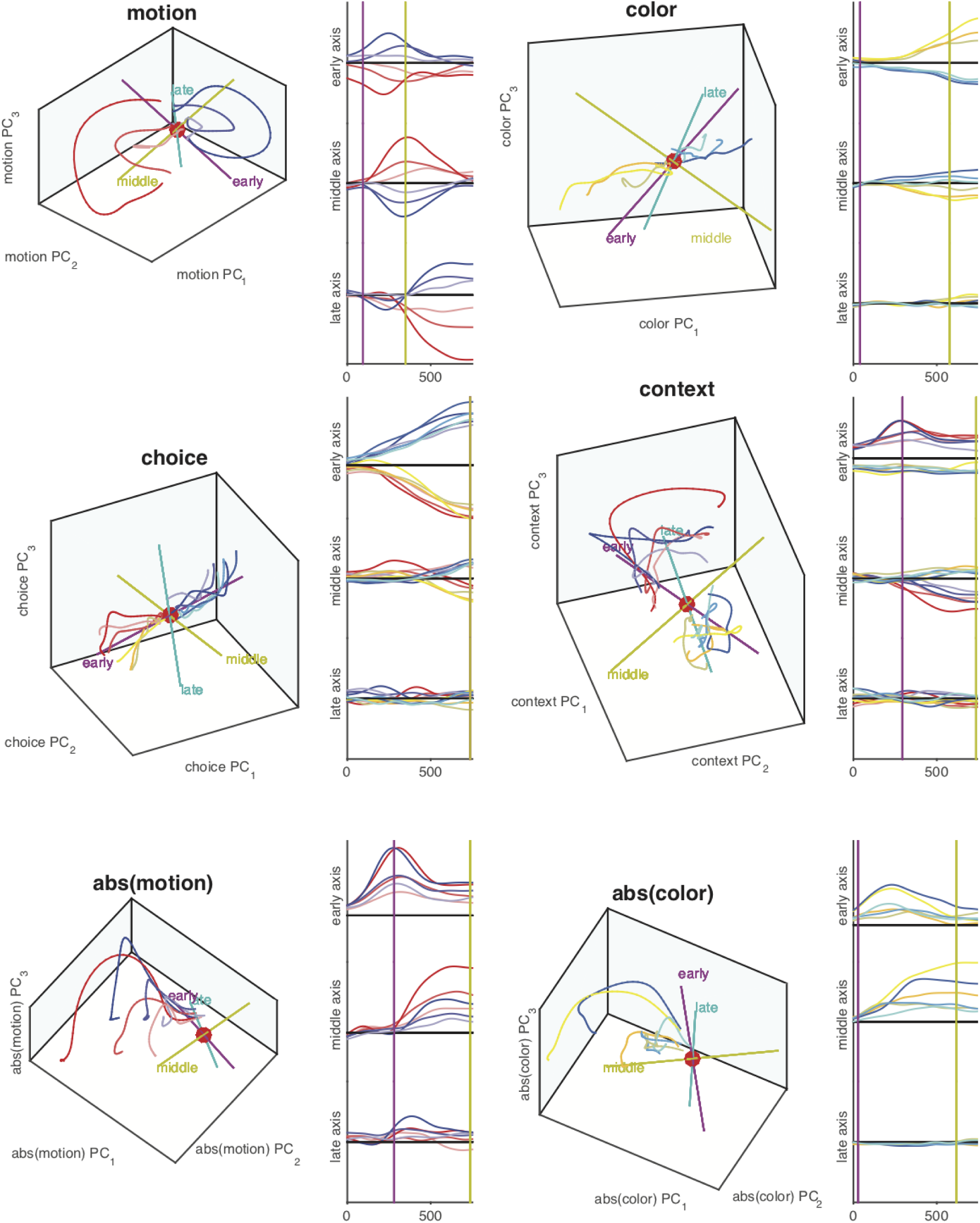
Projections of population PSTH’s for monkey F onto the first, second, and third PC-axes of all task variables subspaces. Plotting conventions and analyses are the same as those for Figure 4. Projected data is averaged over 2-folds of cross validated projections where a random sampling of half of the data was used to estimate parameters and the remaining half used to make projections.

**Figure S12:**
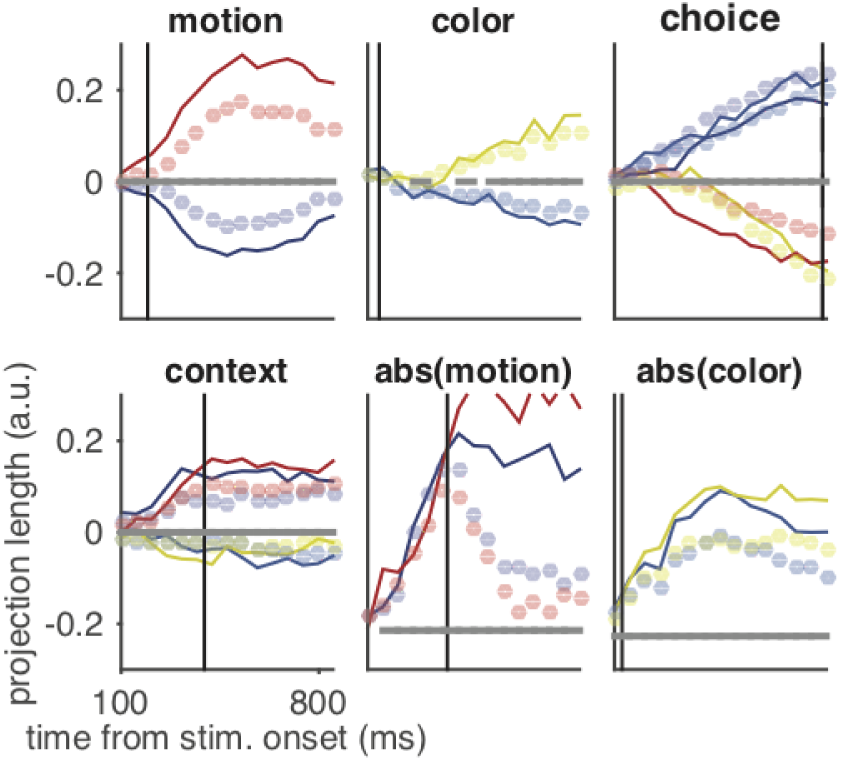
Encoding strength of population pseudosamples for monkey F onto the first three axes of all task variables subspaces. Plotting conventions and analyses are the same as those for Figure 4. Projected data is averaged over 2-folds of cross validated projections where pseudosamples were drawn from held-out trials.

**Figure S13:**
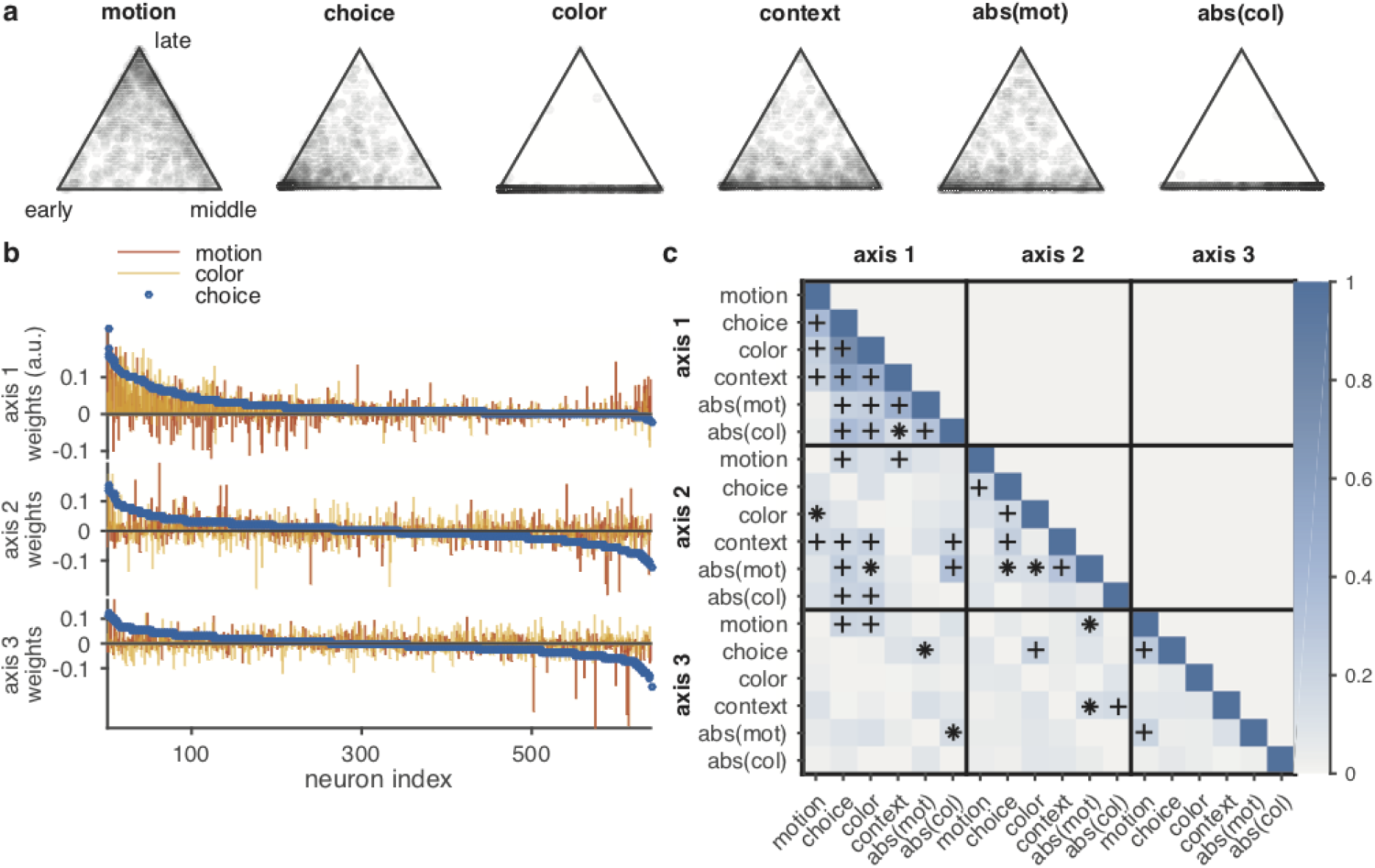
Distribution of variance among seqPCA axes. Monkey F. Plotting conventions are the same as for Figure 5

**Figure S14:**
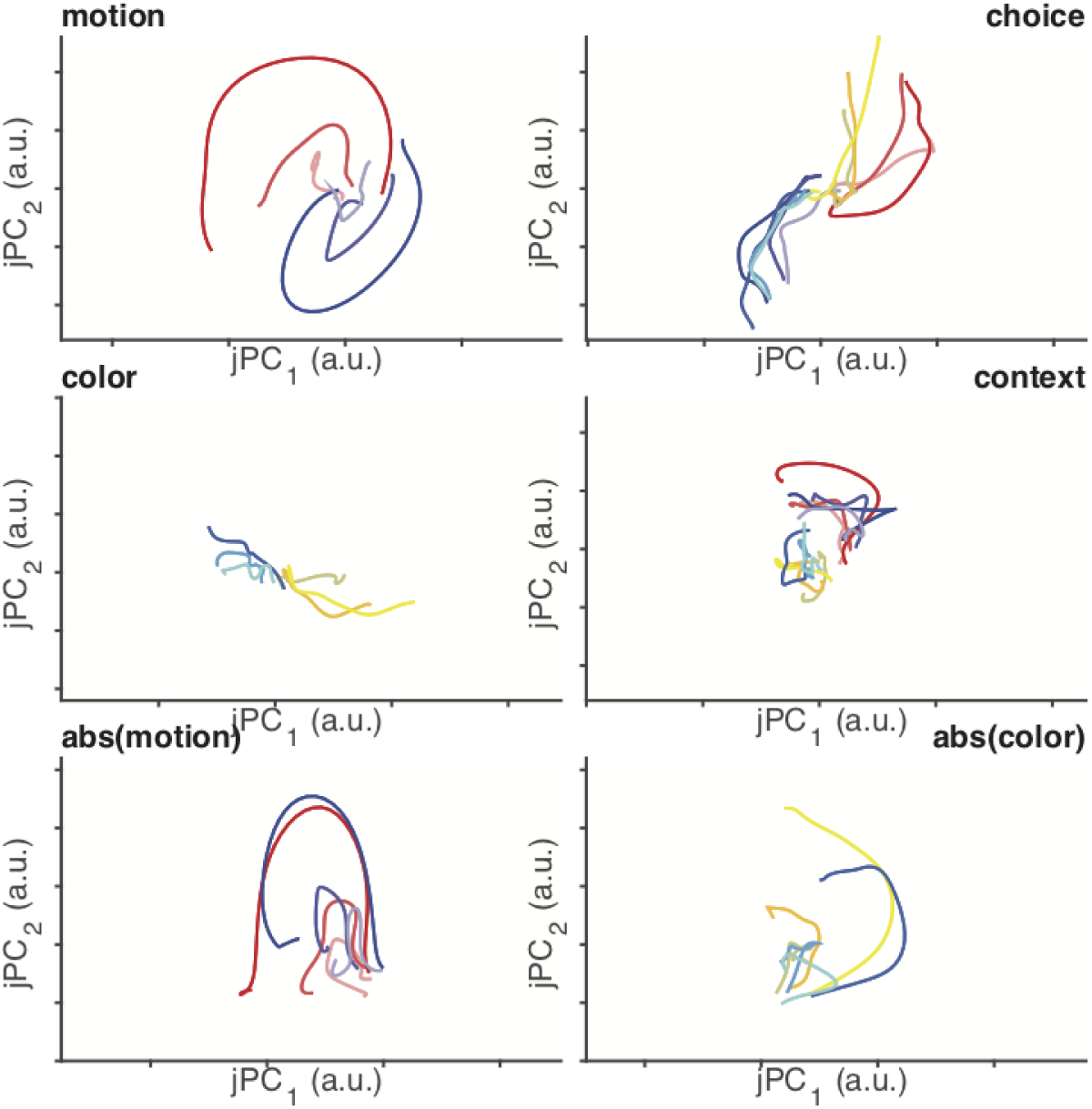
Projections of population PSTH’s for monkey F onto the first, second, and third PC-axes of all task variables subspaces. Plotting conventions and analyses are the same as those for Figure 4. Projected data is averaged over 2-folds of cross validated projections where a random sampling of half of the data was used to estimate parameters and the remaining half used to make projections.

**Figure S15:**
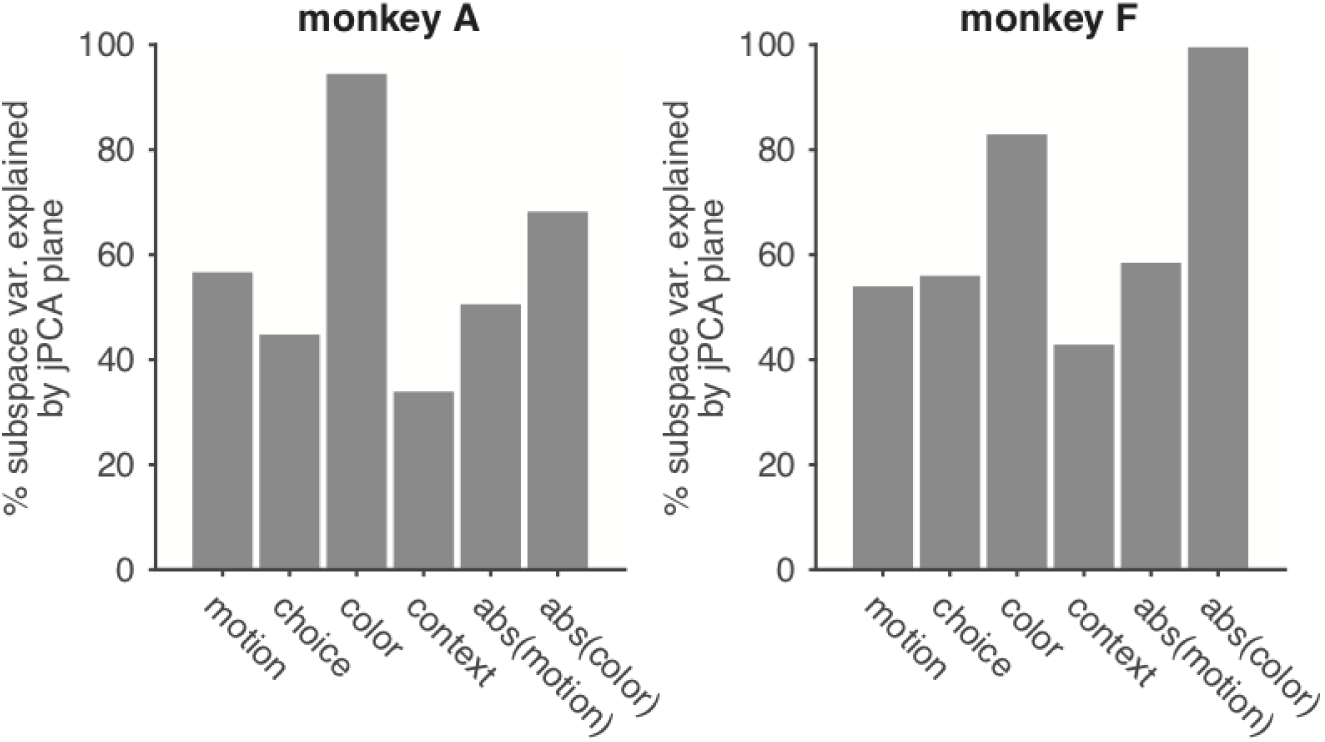
Variance explained by jPCA axes.

**Figure S16:**
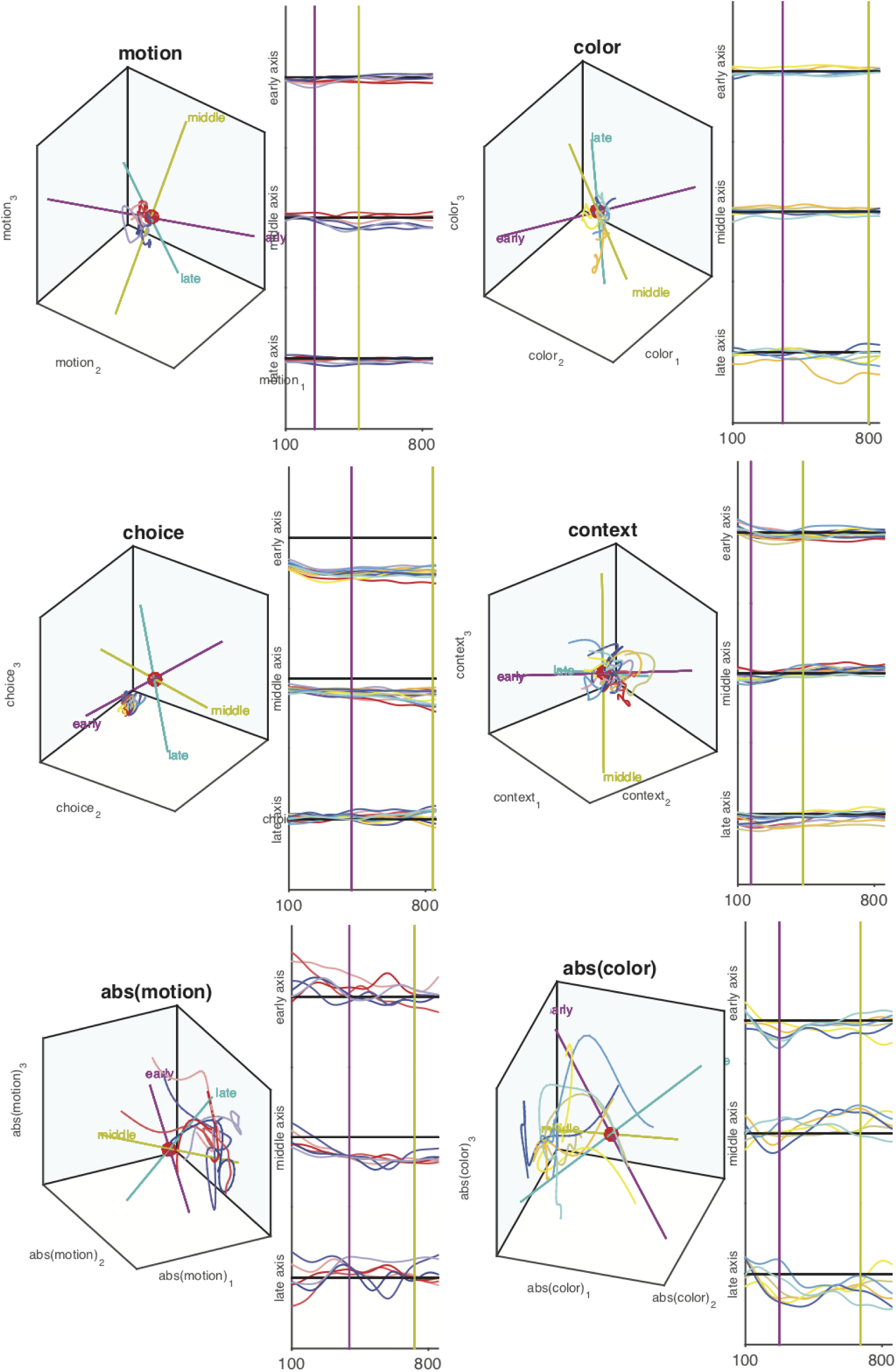
Projections of shuffled population PSTH’s onto task variable subspaces. Monkey A.

**Figure S17:**
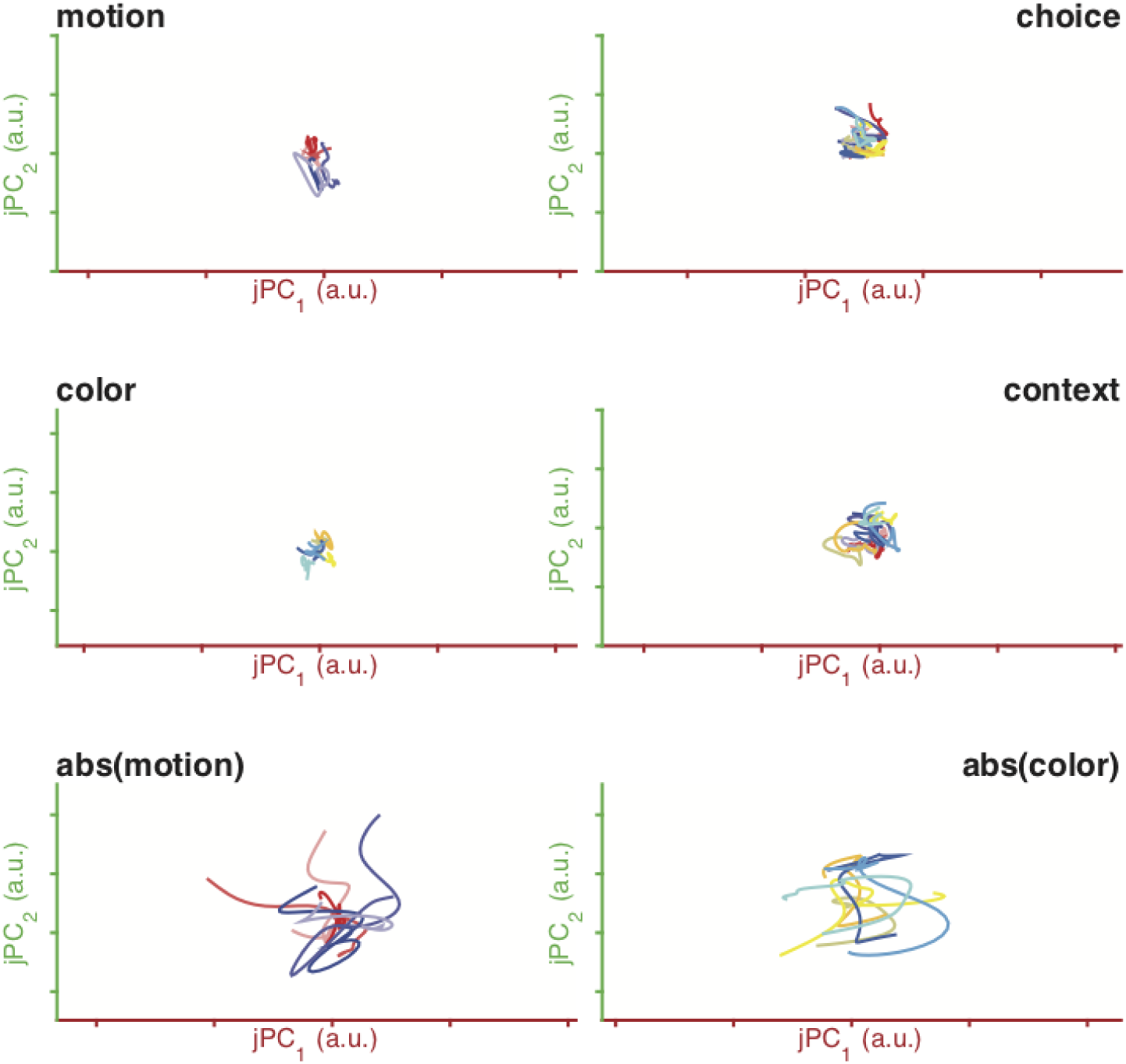
Projections of shuffled population PSTH’s onto jPCA axes. Monkey A.

**Figure S18:**
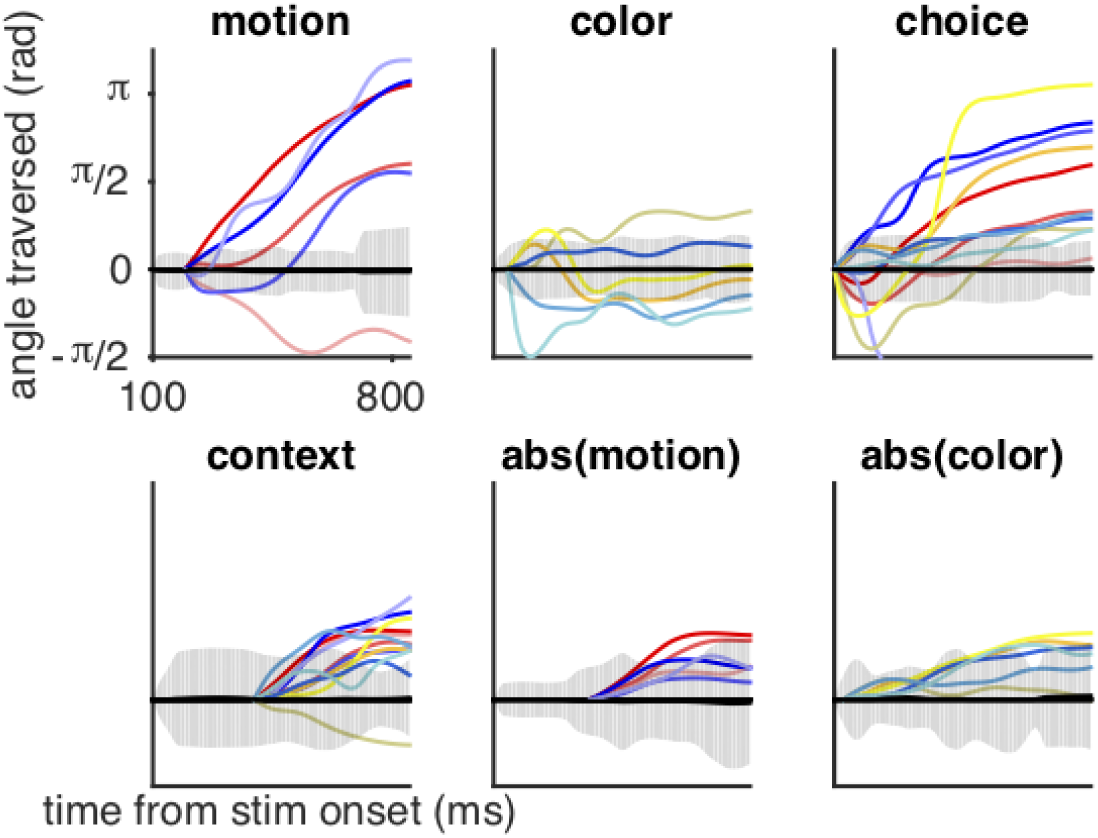
Angle of rotation over time for low-D trajectories of monkey F. Angle of rotation of the low-D trajectories when starting from the start of the middle-axis epoch. Trajectories that are more rotational will appear more monotonic.

**Figure S19:**
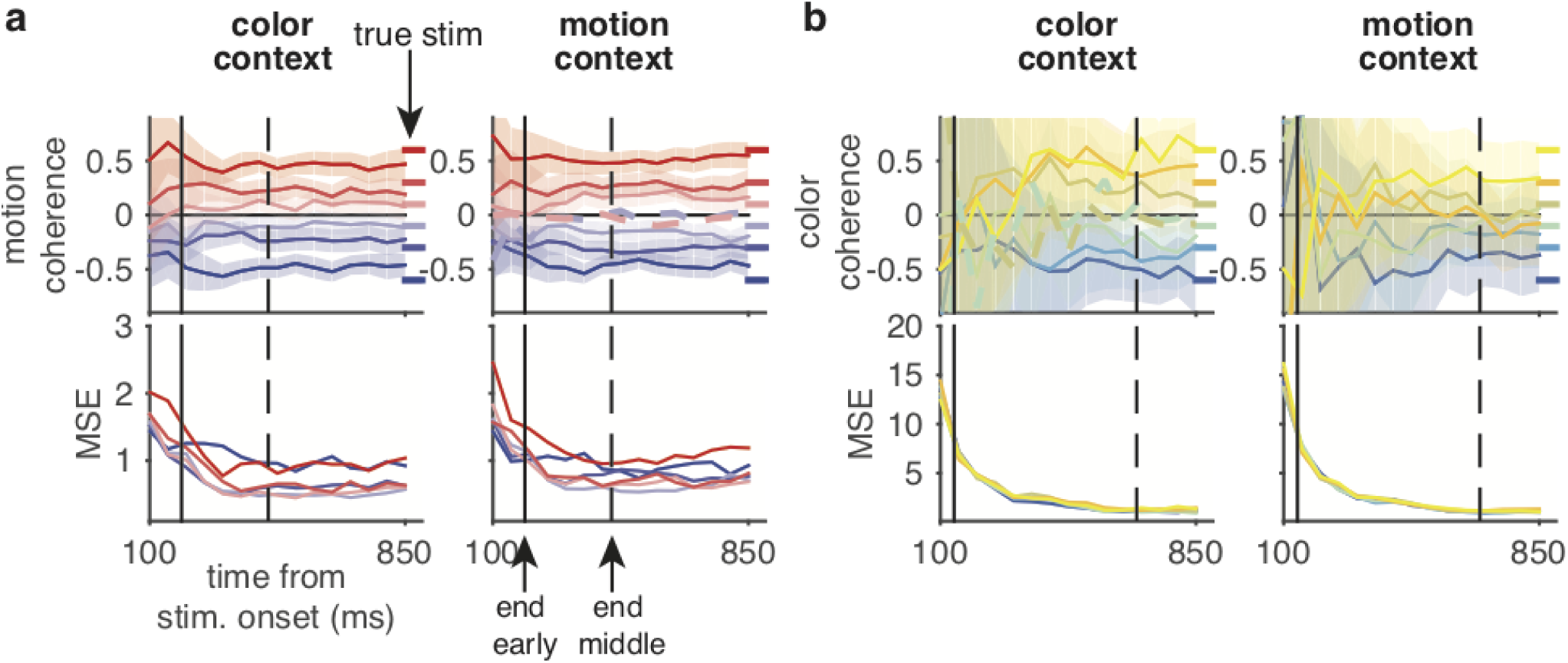
Instantaneous decoding of stimulus for monkey F. Plotting conventions and analyses are the same as for Figure 6

**Figure S20:**
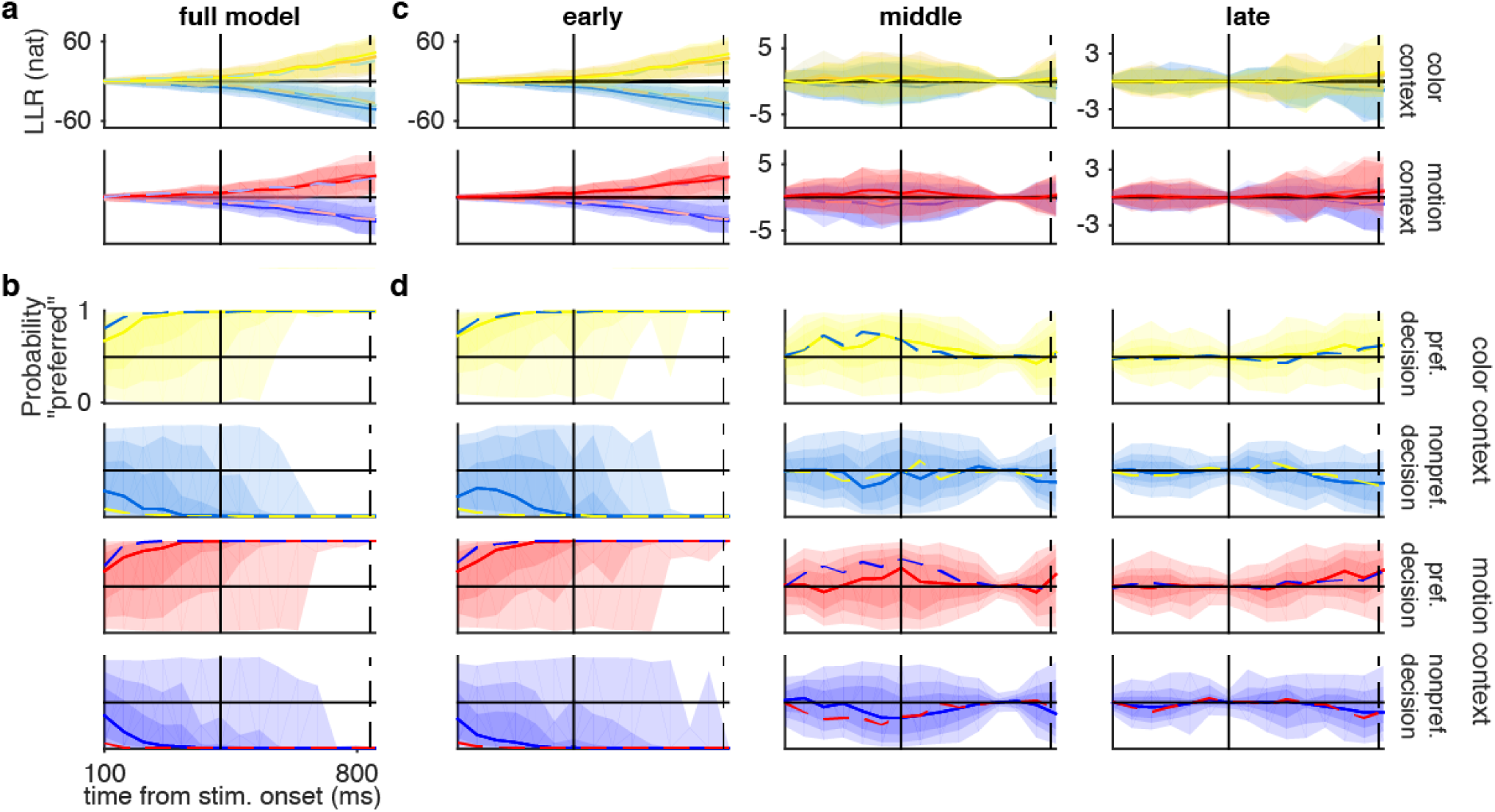
Instantaneous decoding of decision for monkey F. Plotting conventions and analyses are the same as for Figure 6

**Figure S21:**
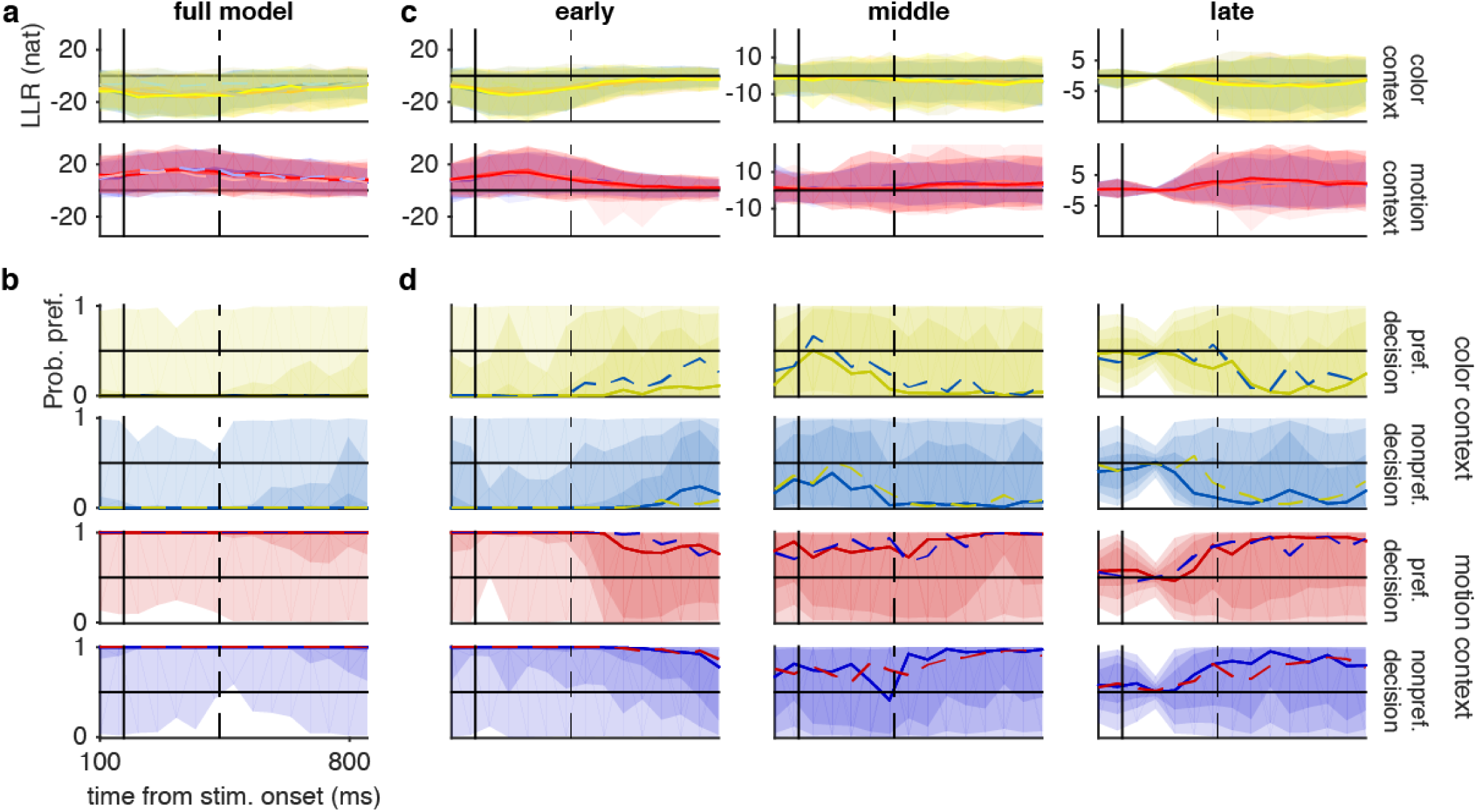
Instantaneous decoding of context for monkey A. **a**) LLRs for monkey A in favor of the motion context using single pseudotrials, sorted by color coherence. Shaded regions indicate 95% quantile intervals for each stimulus strength. Solid lines indicate the median over correct trials. Dashed lines indicate median of error trials. **b**) Probability of the motion context based on corresponding LLRs combined over all stimulus strengths. Solid lines indicate median of correct trials. Dashed lines indicate median of error trials. Shaded regions indicate quantile intervals of correct trials (light-to-dark: 50%, 75%, 95%). Color conventions are the same as in Figure 4. 100 pseudotrials for each of 4-fold cross validation folds used for all analyses.

**Figure S22:**
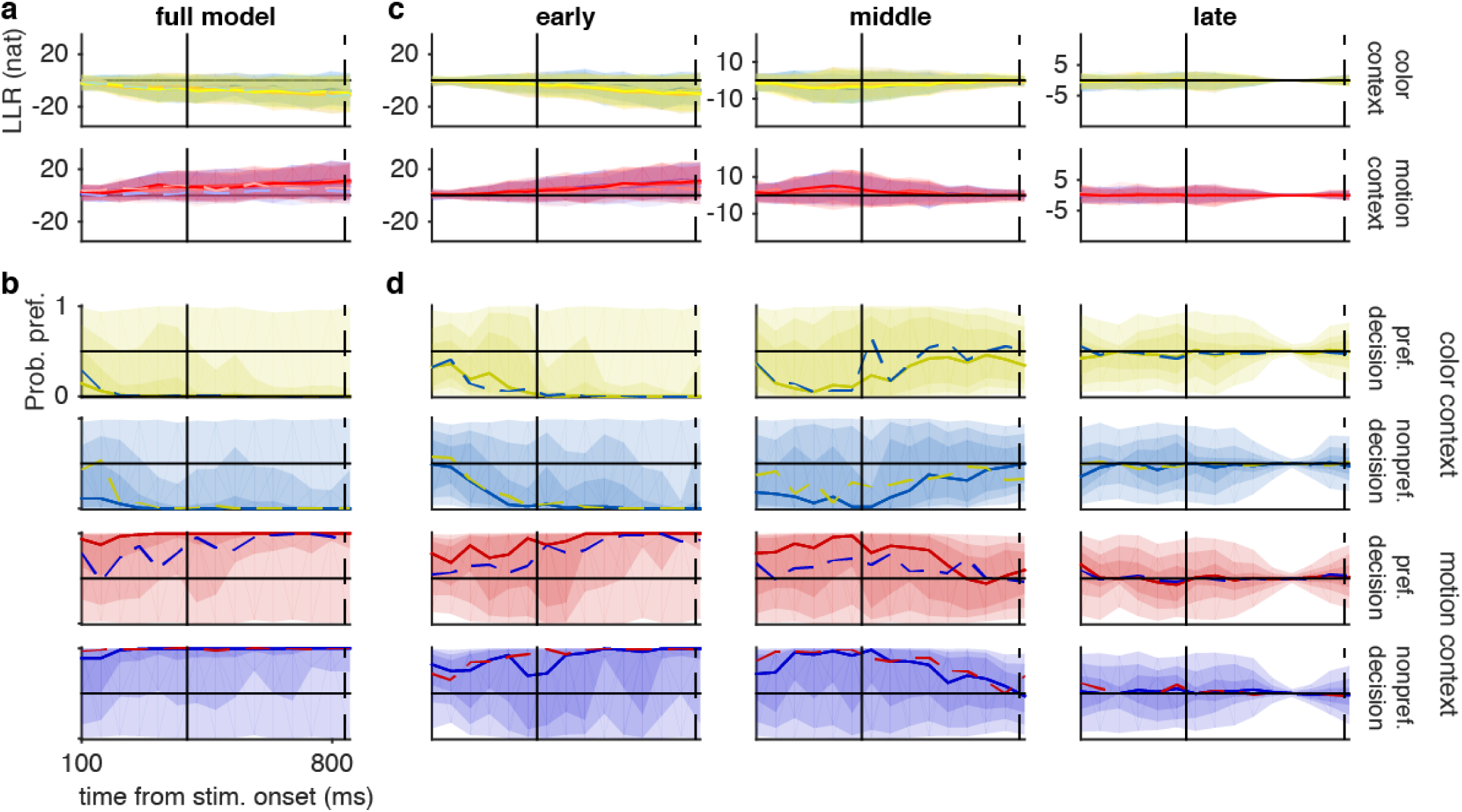
Instantaneous decoding of context for monkey F. Plotting conventions are the same as in Fig. S21. 100 pseudotrials for each of 2-fold cross validation folds used for all analyses.

